# Partial spontaneous intersubunit rotations in actively translating ribosomes

**DOI:** 10.1101/2020.12.23.423865

**Authors:** Tianhan Huang, Junhong Choi, Arjun Prabhakar, Joseph Puglisi, Alexey Petrov

**Affiliations:** Department of Biological Sciences, Auburn University, Auburn AL, USA; Department of Structural Biology, Stanford University School of Medicine, Stanford, CA, USA; Department of Genome Sciences, University of Washington School of Medicine, Seattle, WA, USA; Pacific Biosciences, Menlo Park, CA, USA

**Keywords:** Translation, ribosome, intersubunit rotation, translocation, single molecule FRET

## Abstract

The ribosome is a molecular machine that adopts at least two global states during translation. Two main steps of translation, peptidyl transfer and translocation, are accompanied by counterclockwise and clockwise rotations of the two ribosomal subunits. However, when and why the ribosome alternates between these states remains unclear, with two well supported but conflicting hypotheses. Ribosomes may undergo a single cycle of forward and backward rotations per codon read. Alternatively, in addition to rotations caused by peptidyl transfer and translocation, ribosomes may undergo multiple full spontaneous rotations, with these rotations playing a critical role in elongation and specifically in translocation mechanism. We applied high-speed single-molecule TIRF microscopy to follow translation in real-time. Actively translating ribosomes undergo *partial* spontaneous rotations between three different rotational states. Spontaneous rotations are restricted prior to A-site tRNA decoding. Peptidyl transfer unlocks spontaneous rotations. Consequently, translocation proceeds via a novel rotational state induced by EF-G. Our results bridge both models and provide a coherent view of ribosome dynamics during translation.

## Introduction

The ribosome is a molecular motor that travels directionally along mRNA in three-nucleotide steps as it synthesizes protein. Movement is coupled to changes of the global conformation of the ribosome, which adopts two distinct inter-subunit conformations that differ by an approximately 6-8 degrees rotation of the small (30S) subunit relative to the large (50S) subunit (Fischer et al., 2010; Frank and Agrawal, 2000). The elongation cycle begins with the ribosome in a non-rotated state with an empty A site. tRNA is recruited to the ribosomal A site by elongation factor Tu (EF-Tu); upon cognate tRNA recognition, EF-Tu hydrolyzes GTP and the aa-tRNA is accommodated into the A site of the large subunit, which is rapidly followed by peptidyl transfer. Peptidyl transfer unlocks spontaneous tRNA movement and triggers counterclockwise rotation of the small subunit, thus placing the ribosome in a rotated state. The rotated ribosome is then recognized by EF-G, which catalyzes translocation. During translocation both tRNAs are moved from the A and P sites to the P and E sites correspondingly and the ribosome steps one codon toward the 3’ end of the mRNA. Translocation places the next codon into the now vacant A site and returns the ribosome to the non-rotated state to begin the next elongation cycle (Marshall et al., 2008; Rodnina et al., 1997). Ribosome rotations are essential intermediates of the elongation cycle, as crosslinking subunits by disulfide bonds or adding antibiotics, such as viomycin that block rotations, stalls translation (Aitken and Puglisi, 2010; Horan and Noller, 2007).

Reversible spontaneous ribosomal rotations, where conformational exchange is driven by thermal fluctuations and not by elongation factors, have been observed with preformed elongation complexes (Cornish et al., 2008; Sharma et al., 2016) using single-molecule FRET between fluorophores attached to the S6 protein of the 30S subunit and the L9 protein of the 50S subunit (Ermolenko et al., 2007). Spontaneous rotations were found to be infrequent in ribosomes with peptidyl-tRNA in P site. Upon A-site tRNA binding, peptidyl transfer unlocks spontaneous rotations in the pre-translocation state, allowing ribosomes to fluctuate between non-rotated and rotated states. Ensemble experiments indicated that EF-G biases fluctuating ribosomes into a rotated state, and then catalyzes translocation during which ribosomes transition into a non-rotated state. These results led to a model where the ribosome exploits full reversible spontaneous rotations that are coupled to tRNA spontaneous movements to promote translocation. In this model ribosome is a Brownian motor with low energy barriers between rotational states that could be overcome by thermal energy. Factors capture these fluctuations and recuperating energy is provided by the GTP hydrolysis and peptide bond formation (Cornish et al., 2008; Noller et al., 2017).

Following actively translating ribosomes by single-molecule methods resulted in a different model. These experiments employed rRNA modified ribosomes carrying fluorophores on DNA oligomers bound to extensions of helix 44 (h44) of 30S subunit and helix 101 (h101) of 50S subunit (Dorywalska et al., 2005). Similarly, ribosomes occupied non-rotated state prior to A-site tRNA binding and rotated state before translocation. However, no spontaneous rotations were observed in either state. Thus, the functional states of the ribosome would be separated by larger, bigger than RT, energy barriers. Peptidyl transfer and GTP hydrolysis by EF-G provide the energy that drives conformational change of the ribosome as well as movement of the ribosome over mRNA during translocation. The predictable change of the ribosomal conformation during elongation enabled tracking of individual ribosomes as they translated multiple codons (Aitken and Puglisi, 2010). This provided a basis for deciphering mechanisms of elongation (Chen et al., 2013; Choi and Puglisi, 2017; Navon et al., 2016), decoding events(Chen et al., 2015; Tsai et al., 2014), and translation acting antibiotics(Johansson et al., 2014; Tsai et al., 2013).

Both models are well supported experimentally and the dissonance between them precluded building a consistent model of elongation and understanding how ribosome moves over mRNA. Here we used both S6 - L9 and h44 - h101 FRET to follow conformation of actively translating ribosomes and showed their full equivalence. Furthermore, using highspeed single molecule imaging, we observed transient rotational intermediates during translation. Our results bridge both models and provide a coherent view of inter-subunit dynamics during translation.

## Results

### Real-time tracking of 70S formation with S6-L9 ribosomes

We started by monitoring in real time intersubunit rotation in translating ribosomes using the S6-L9 FRET signal. 70S ribosome formation could be followed by appearance of FRET between dyes on S6 and L9. The subsequent intersubunit rotations change the distance between S6 and L9 proteins, which will manifest as a change in FRET efficiency. The experiments were performed using same experimental conditions and conventional TIRF single-molecule microscopy at 200 ms time resolution that were previously used to follow translation with h44-h101 ribosomes.

Joining 50S subunits to 30S preinitiation complexes finalizes initiation and results in elongation-competent ribosomes. The initiation complexes containing 30S-Cy3 ribosomal subunits, mRNA, fMet-tRNA^fMet^ and IF2 were immobilized on the surface of the microscope slide. Translation was started by delivery of Cy5-labeled 50S ribosomal subunits simultaneously with the start of imaging, thus allowing real-time observation of translation (Supplementary Figure 1A). Subunit joining was fast at 25 nM of 50S subunits with *k*_obs_ being 0.13±0.009 s^−1^. Kinetics of 50S subunit arrival was best described by a single-exponential fit indicating functional uniformity of initiation complexes. The 50S subunit arrival time was concentration dependent, as expected for a bimolecular reaction. The 2^nd^ order rate constant was calculated as the slope of the concentration dependence curve, being 3.8 μM^−1^s^−1^ (Supplementary Figure 1C; Supplementary Table 1). It closely matches the 4.1 μM^−1^s^−1^ rate constant from previous measurements with h44-h101 ribosomes (Marshall et al., 2008, 2009) and the 9 μM^−1^s^−1^ subunit joining rate measured with S6-L9 ribosomes by singlemolecule experiments (Ling and Ermolenko, 2015). Prior structural and single-molecule experiments observed that subunits initially join in the rotated or semi-rotated conformation. Rapid GTP hydrolysis by IF2 triggers reverse rotation of the ribosomal subunits (Kaledhonkar et al., 2019; Ling and Ermolenko, 2015; Sprink et al., 2016). The pre-rotation state is very transient and thus seldomly observed in single-molecule traces. Consistently, subunit joining puts ribosomes into a high FRET state with efficiency of 0.6 (Supplementary Figure 1B) that corresponds to a non-rotated conformation of the ribosome. Spontaneous, i.e. thermally driven, rotations were infrequent with 92% of ribosomes never leaving the high FRET state. This fully agrees with previous ensemble and single-molecule measurements with h44-h101 and S6-L9 ribosomes that demonstrated no or very slow spontaneous ribosomal rotation in 70S ribosomes with either fMet-tRNA^fMet^ or peptidyl-tRNA bound to the P-site (Cornish et al., 2008; Marshall et al., 2009). Together it validates our experimental setup and readies us to address elongation dynamics.

### tRNA binding changes ribosome conformation with S6-L9 ribosomes

Joining of 50S subunit and A-site tRNA arrival were followed by supplementing delivery mix with A-site ternary complex (TC). Upon peptide bond formation, the ribosome adopts a new global state in which it solely or predominantly occupies a rotated conformation (Cornish et al., 2008; Ermolenko et al., 2007; Marshall et al., 2008). Forward intersubunit rotation increases the distance between S6 and L9 proteins that manifests as a decrease in FRET. In these experiments, upon subunit joining, ribosomes rapidly shifted into a lower FRET state (Figure 1A). The transition rate is best approximated by a single-exponential fit, with a rapid reaction rate of 0.208±0.013 s^−1^ at 25 nM ternary complex. The transition rate was dependent on the ternary complex concentration. Rate constant was determined as the slope of concentration dependence curve, being 7.5 μM^−1^s^−1^. This is similar to A-site tRNA binding rates that were found to be between 1.1-3.7 μM^−1^s^−1^ in previous single-molecule experiments (Supplementary Table 1). Importantly, peptidyl transfer occurs on millisecond to a tens of milliseconds time scale (Johansson et al., 2008; Katunin et al., 2002) and thus does not contribute to the measured rate, as it is too fast to be detected. Thus, this conformational transition is caused by tRNA binding which is followed by rapid peptidyl transfer.

**Figure 1.**
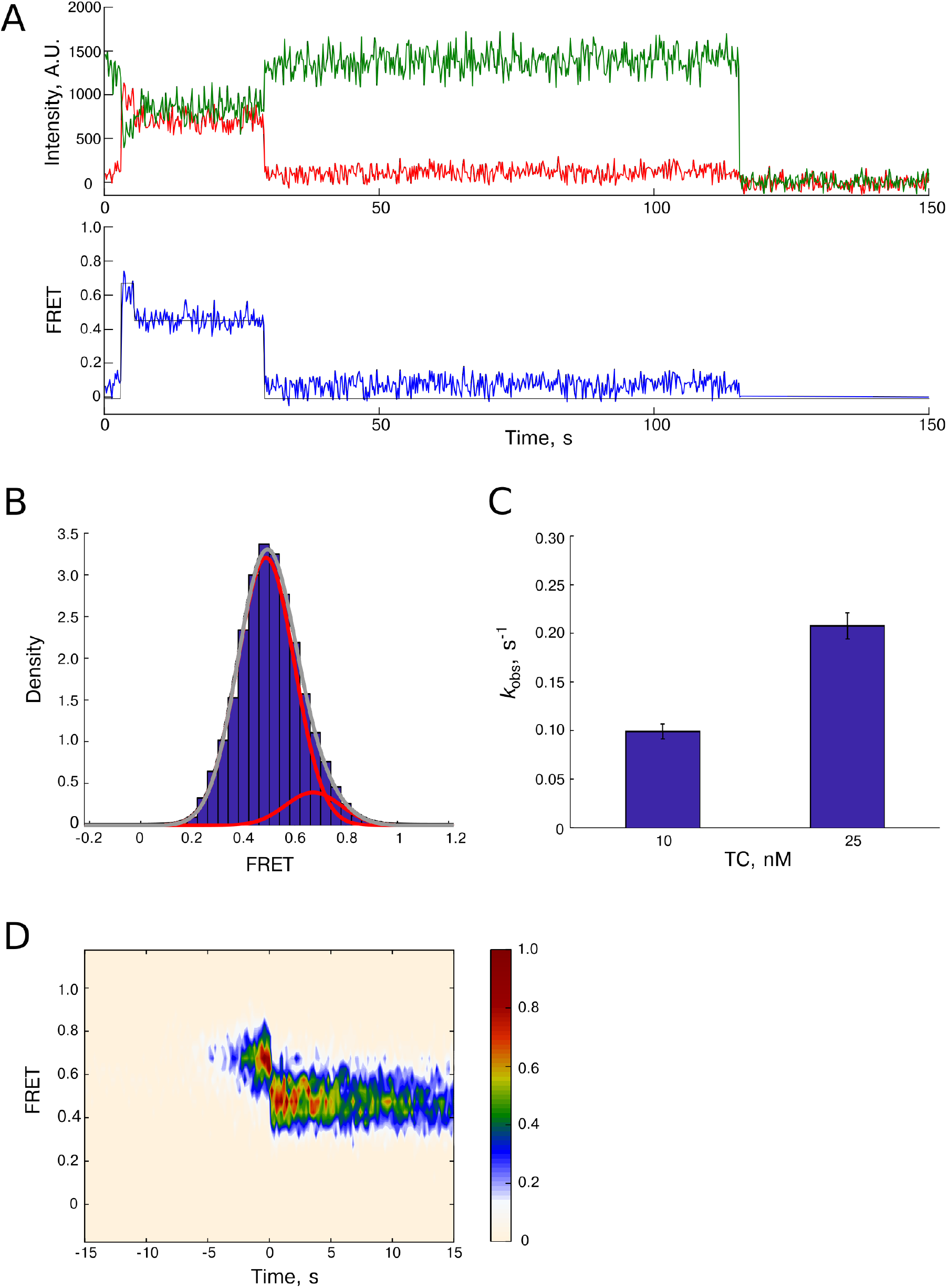
Ribosome conformational dynamics upon tRNA binding. This experiment was performed using S6-L9 labeled ribosomes. 30S initiation complexes comprised of 30S-Cy3 ribosomal subunits, initiator tRNA, IF2 and GTP were assembled over biotinylated (FK)6 mRNA. Complexes were immobilized on the surface and 50S-Cy5 ribosomal subunits, ternary complex of EF-Tu, Phe-tRNA^Phe^ and GTP were delivered concurrently with the start of imaging. **A.** Example trace. 30S-Cy3 fluorescence is shown in green and 50S-Cy5 fluorescence is shown in red. 70S ribosome is formed rapidly at ~ 3 s time point. Promptly, after the ~5 s mark, the ribosome rotates resulting in a FRET decrease. FRET intensity remains constant until Cy5 photobleaching at 30 s. The FRET idealization used in state analysis is shown by a hairline overlaying FRET values in the bottom panel. **B.** FRET intensity histogram shows the appearance of the major low FRET peak. FRET decrease is due to increased distance between dyes, which corresponds to the ribosome adopting a rotated conformation. The smaller ~0.65 FRET peak corresponds to the transient non-rotated state before tRNA binding, which corresponds to the 70S ribosomes with fMet-tRNA^fMet^ in P-site. N = 107. **C.** The timing of the ribosome rotation was dependent on ternary complex concentration. N = 154, 412, and 80. Error bars are errors of the fit. The reaction rate determined as a slope of linear regression was found to be 6.4 μM^−1^s^−1^. **D.** Post-sync plot demonstrates lack of detectable spontaneous rotations at these experimental conditions. Traces were synchronized at the high-to-low FRET transition. Color shows experimental probability of observing FRET value.

tRNA binding results in pre-translocation ribosomes. Most ribosomes showed a single shift from high to lower FRET state that corresponded to tRNA binding. Virtually no additional FRET changes that would correspond to spontaneous rotations were observed (Supplementary Figure 2B). This is further corroborated by a post-sync plot of FRET intensities before and after tRNA arrival that show ribosomes predominantly occupy two distinct states separated by tRNA binding (Figure 1D). Accordingly, the overall FRET intensity histogram was best approximated by a double Gaussian fit (Figure 1B). The minor peak (high FRET) corresponds to the transient non-rotated ribosomes before tRNA binding. The major peak (lower FRET) corresponds to the ribosomes continuously residing in the rotated state after tRNA binding. The FRET intensity histograms of individual states were single-modal Gaussians (Supplementary Figure 3). These results are congruent with previous measurements of actively translating h44-h101 ribosomes that demonstrated single FRET transition upon tRNA binding.

### EF-G catalyzes reverse ribosomal rotation in actively translating S6-L9 ribosomes

To examine ribosome conformation during translocation, we delivered a mixture of 50S subunits, A-site TC and EF-G to immobilized initiation complexes. Upon subunit joining, A-site tRNA binding promoted forward rotation, which was detected by a decrease in FRET. In the presence of EF-G, ribosomes transitioned back to the high FRET state, which corresponds to reverse rotation of the ribosomes during translocation.

The kinetic of reverse rotation was single exponential and dependent on EF-G concentration. The rate constant was determined as the slope of the reaction rate vs EF-G concentration plot and was found to be 4.4 μM^−1^s^−1^. Similarly to previous results (Aitken and Puglisi, 2010; Chen et al., 2012; Marshall et al., 2008) most molecules did not exhibit spontaneous rotations (Supplementary Figure 2C).

Spontaneous transitions are rare (9% in 70S initiation complexes, 7% in pre-translocation complexes, and 11% in post-translocation complexes waiting for the second tRNA) and slow with rates of 0.01-0.1 s^−1^. Consequently, each functional state of the ribosome is characterized by a single-modal FRET distribution, corresponding to either high or low FRET (Supplementary Figure 3), and the overall FRET distribution is bimodal (Figure 2B). Thus, the ribosome undergoes forward rotation upon tRNA binding followed by peptide bond formation and reverse rotation upon translocation, with no observable spontaneous rotations on this timescale.

**Figure 2.**
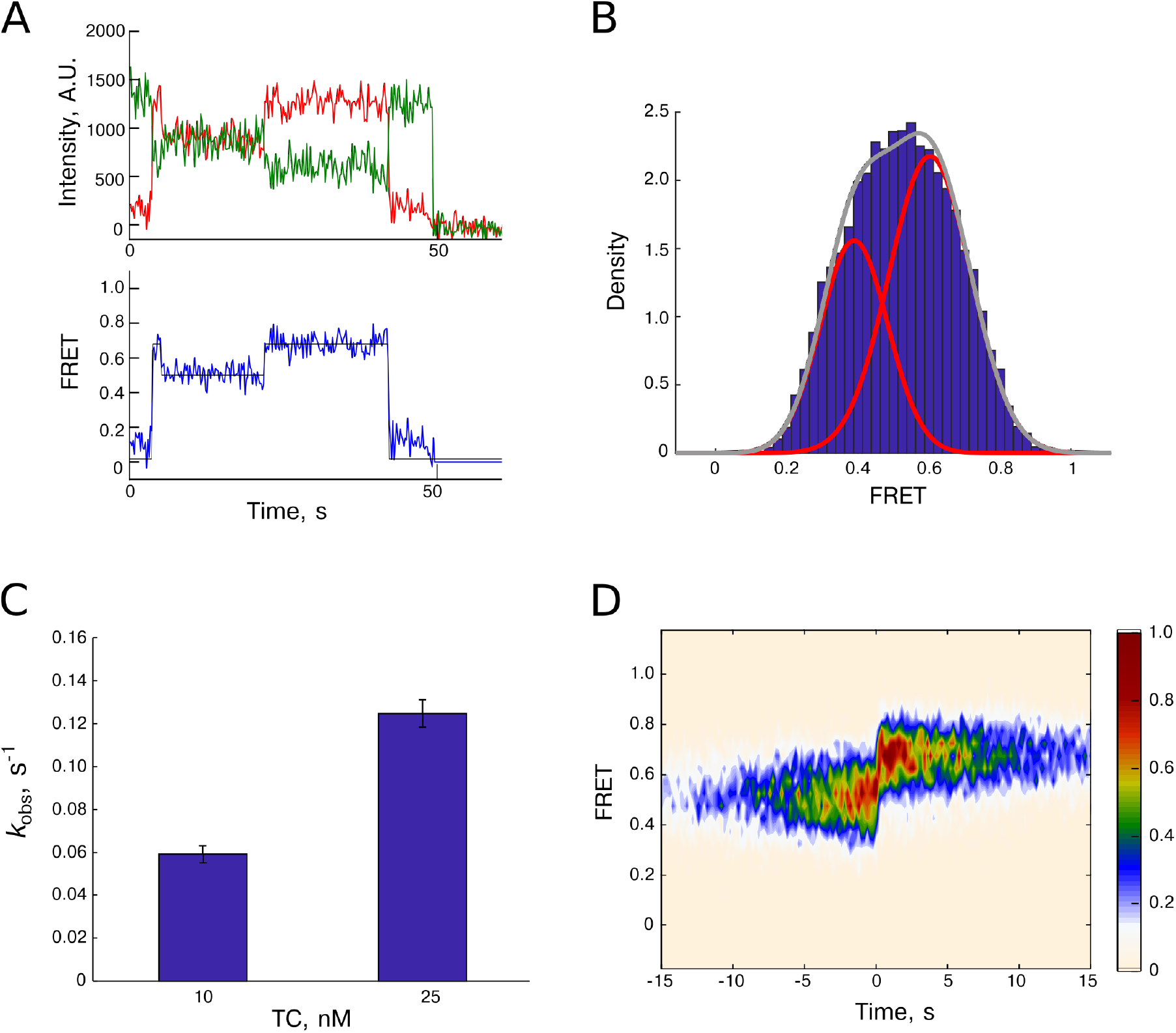
Following the ribosome through the first elongation cycle. 30S initiation complexes comprised of 30S-S6-Cy3 ribosomal subunits, initiator tRNA, and IF2 were assembled over biotinylated (FK)6 mRNA. Complexes were immobilized on the surface. 50S-L9-Cy5 ribosomal subunits and ternary complex of EF-Tu, Phe-tRNA^Phe^ and GTP and EF-G were delivered concurrently with the start of imaging. **A.** Example trace. 70S ribosome is formed rapidly at ~ 3 s time point, which is followed by tRNA binding that manifested as a decrease in FRET. At ~20 s ribosome undergoes reverse rotation caused by EF-G and returns to the non-rotated conformation. **B.** FRET intensity histogram is bimodal, with the low FRET peak corresponding to pre-translocation ribosomes and the high FRET peak corresponding to ribosomes before tRNA binding and after translocation. **C.** The rate of reverse ribosome rotation was dependent on EF-G concentration. Error bars are errors of the fit. The reaction rate was determined as a slope of linear regression and found to be 4.4 μM^−1^s^−1^. **D.** Post-sync plot demonstrates lack of spontaneous fluctuations at these experimental conditions.

### Continuous translation over multiple codons

Can predictable alternations between high and lower FRET states in response to tRNA binding and translocation be used to follow translation over multiple rounds of elongation with S6-L9 ribosomes? To explore this possibility, a mRNA encoding an FKF peptide was translated in the presence of Phe-tRNA^Phe^ and Lys-tRNA^Lys^. Changes in FRET were used to identify tRNA binding and translocation with each cycle of high-to-low-to-high FRET change corresponding to translation of one codon. Translation of up to three codons was observed (Supplementary Figure 4B). Replacing the mRNA with a longer, 12-codon (FK)_6_ mRNA, increased the number of observed rotations, with robust translation seen up to 12 codons (Supplementary Figure 4D). In control experiment where the second codon tRNA (Lys-tRNA^Lys^) was omitted, the majority (87%) of the ribosomes translated only a single codon (Supplementary Figure 4C). Thus, FRET transitions could be used to follow mRNA translation.

Previous single-molecule experiments with ribosomes labeled on the rRNA demonstrated that reverse (clockwise) rotation coincides with EF-G occupancy of the ribosome (Chen et al., 2012). This observation served as a basis to establish mechanisms of elongation and translational recoding events (Chen et al., 2012; Choi and Puglisi, 2017; Prabhakar et al., 2017). To see if reverse rotations are directly promoted by EF-G in S6-L9 ribosomes, we directly examined the relationship between ribosome occupancy by EF-G and ribosomal conformation. We labeled EF-G with Alexa588 dye as previously described (Chen et al., 2013), and used EF-G-Alexa588 in real-time translation experiments with (FK)_6_ mRNA. Translation was followed with S6-L9 FRET and EF-G binding was detected by direct dye illumination. To observe simultaneously intersubunit FRET and directly illuminated EF-G, we employed ZMW single-molecule microscopy (Chen et al., 2014). As expected, ribosomes translated mRNA over multiple codons (Figure 3A), with 49% of the low to high FRET transitions coinciding with EF-G binding events (Figure 3B). Due to the long exposure time (100 ms), the EF-G dwell time behaved single exponentially with dissociation rate *k*_obs_ = 14 s^−1^ (Figure 3C). This corresponds to the EF-G dwell time of ~ 71 ms, which is in agreement with the previously reported 60 ms dwell time under the same experimental conditions (Chen et al., 2013). The short EF-G dwell time suggests that during the remaining 51% of transitions, EF-G was not detected due to translocation occurring faster than 100 ms exposure time. Next, we synchronized single-molecule trajectories to the low-to-high FRET transition. The strong overlap between ribosome rotation and EF-G residency indicates that reverse intersubunit rotation is promoted by EF-G (Figure 3B).

**Figure 3.**
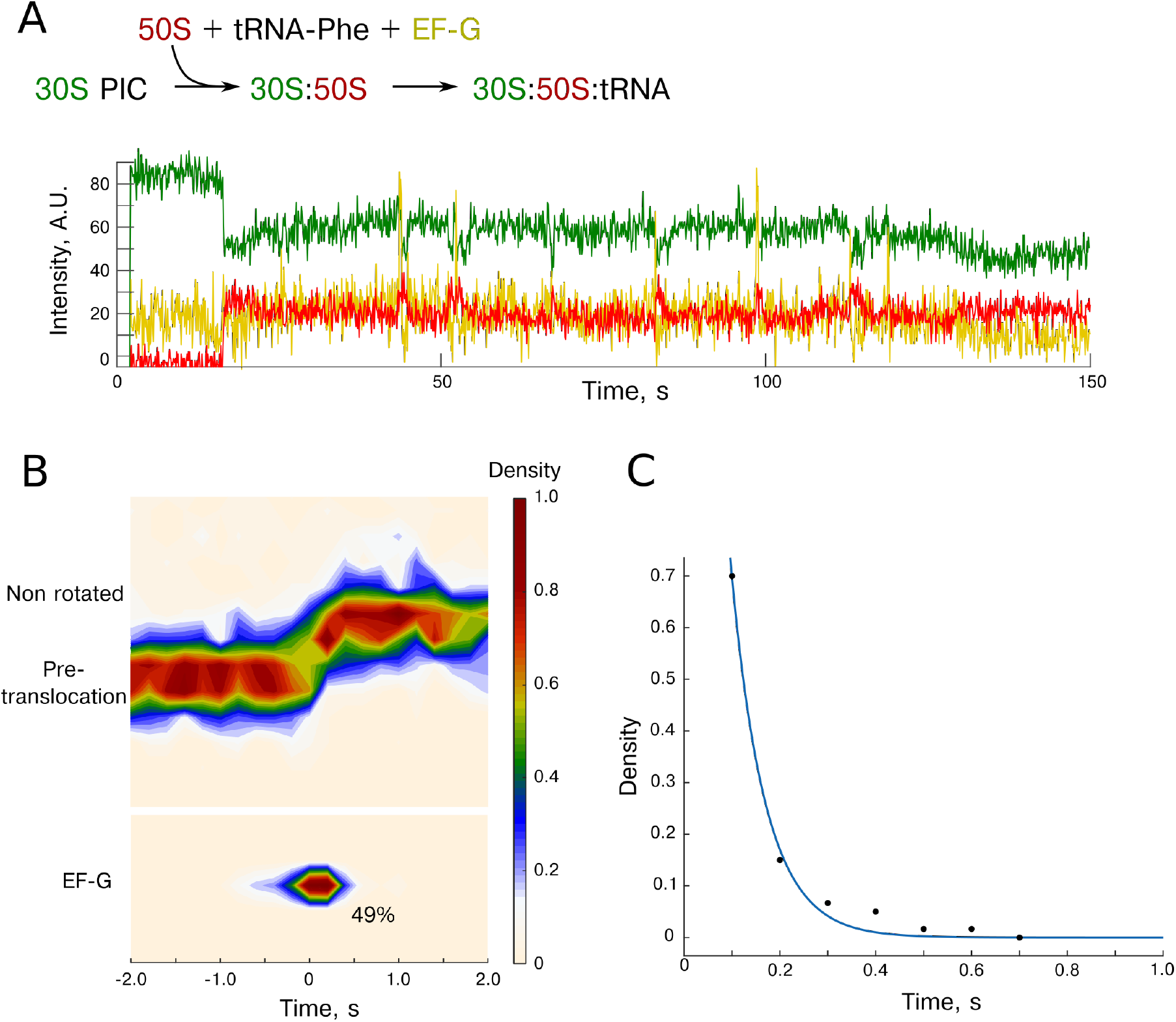
Correlating EF-G binding and ribosome reverse rotation. Translation of the (FK)6 mRNA in the presence of Phe-tRNA^Phe^, Lys-tRNA^Lys^, and EF-G-Alexa588. The EF-G fluorescence is shown in yellow. **A.** Example trace. The ribosome changes between rotated and non-rotated states as it translates mRNA. The EF-G occupancy coincides with reverse ribosome rotation. **B.** Traces were synchronized to the reverse rotation. The FRET transitions (top panel) coincide with EF-G occupancy (bottom panel). **C.** EF-G dwell times.

These results show full agreement between intersubunit rotations in actively translating S6-L9 ribosomes and plethora of single-molecule experiments that utilized h44-h101 FRET to follow translation. Upon subunit joining the ribosomes occupy a non-rotated state characterized by high FRET. Next, tRNA binding promotes transition into a lower FRET state. Translocation returns the ribosome into the non-rotated conformation and this conformational transition coincides with EF-G occupancy of the ribosome. This clockwork like exchange of states allows reliable tracking of translating ribosomes over multiple codons.

### High time-resolution imaging of translation

The described results provide coherent description of ribosome rotations using two distinct FRET pairs in actively translating ribosomes. However, these results do not explain presence of spontaneous rotations observed with S6-L9 ribosomes in static experiments that are also strongly supported experimentally (Belardinelli et al., 2016; Cornish et al., 2008; Sharma et al., 2016). The ensemble model of translocation also implies presence of fast ribosomal rotations (Belardinelli et al., 2016; Sharma et al., 2016). We thus hypothesized that in actively translating ribosomes spontaneous rotations are too fast to be detected. If spontaneous rotations exist, they must be faster than the time resolution of the described experiments; i.e., under 200-400 ms. Most of these fast events would be unobservable. However, some of them might take longer due to the innate stochasticity of chemical reactions and thus would be detectable, therefore potentially contributing to occasionally observed spontaneous rotations.

To determine existence of rapid spontaneous rotations, we constructed a fast timescale, prism-based total internal reflection microscope capable of single molecule imaging at 10 ms resolution (Supplementary Figure 5). High-speed imaging requires an increase in the illumination power. Consequently, S6-L9 ribosomes are not suitable for high-speed measurements due to the low photostability of protein-conjugated fluorophores, which leads to rapid photobleaching. Based on the established full equivalence of the S6-L9 and h44-h101 signals, we decided to use an h44-h101 FRET pair due to its higher photostability. First, we repeated the experiments described above under low-speed conditions and found no differences from S6-L9 ribosomes or from previous h44-h101 results (Supplementary Figure 6).

Next, we observed translation at 10 ms resolution from initiation through elongation, beginning with subunit joining. High temporal resolution allowed reliable observation of the transient initiation steps. 75 percent of the ribosomes joined in the intermediate FRET state and then rapidly transitioned into a high FRET state (Supplementary Figure 7A). The initial FRET state corresponds to the 70S initiation complexes with IF2 and GTP. GTP hydrolysis by IF2 triggers clockwise ribosomal rotation, resulting in non-rotated ribosomes characterized by high FRET. 70S-IF2-GTP ribosomes occupy a unique intermediate rotational state that is distinct from states observed during elongation. It sits between the non-rotated ribosomes and spontaneously rotated ribosomes described below (Supplementary Figure 7B). These results are in agreement with time-resolved Cryo-EM that found that IF2 dissociation resulted in a partial 3 degrees reverse rotation (Kaledhonkar et al., 2019), and single-molecule experiments (Ling and Ermolenko, 2015). Thus, ribosomes join in an intermediate rotational state, which we are going to call the first intermediate state.

Subunit joining puts ribosomes into a high FRET state with slow spontaneous transitions into the lower FRET state. This transition is characterized by ~0.1 decrease in FRET efficiency (Supplementary Table 3). Excursions into the lower FRET state are common, with 24% of ribosomes showing them (Supplementary Figure 7B). Ribosomes spent most of the time in the high FRET state and rapidly reverted from lower FRET to high FRET. Statistical analysis of dwell times reflects this behavior, with high FRET mean dwell time being 29.0 s (*k_obs_* = 0.035 s^−1^) and the low FRET mean dwell time being 0.8 s (*k_obs_* = 1.2s^−1^) correspondingly (Supplementary Table 2A). Dwell times of both states are single exponential suggesting a simple one-step mechanism underlying spontaneous rotations. Therefore, 70S ribosomes can spontaneously transition between two rotational states, but highly favor the non-rotated state.

Next, we added A-site tRNA into the delivery mix. Same as above, ribosomes joined in the high 0.63 FRET state. tRNA binding promoted transition into a pre-translocation state characterized by long-lived lower 0.53 FRET (Figure 4, Supplementary Figure 9). We used this transition to identify active 70S complexes, to eliminate the possibility that ribosomes fluctuating prior tRNA binding were inactive or represented off-pathway states. The frequency of spontaneous rotations in active ribosomes prior to tRNA binding (24%) was the same as in the overall ribosomal pool in the previous experiment (24%). Thus, spontaneous rotations of 70S complexes prior to A-site tRNA recruitment are rare but are part of normal translation. Importantly, high-speed imaging revealed that pre-translocation ribosomes also underwent spontaneous transitions into the high-FRET state as well as into a new, previously unobserved, 0.48 low-FRET state (Figure 4A, Supplementary Figure 8A and Supplementary Table 3). Spontaneous rotations were common, with 39% of ribosomes undergoing transient excursions into a non-rotated state and 51% showing spontaneous transitions into a novel low-FRET state. In total, 71% of ribosomes exhibited spontaneous rotations in the pre-translocation state. The three rotational conformations are distinct, as indicated by overlaid FRET intensity histograms of individual states (Figure 4B). For clarity we are going to reference them as high, first intermediate, second intermediate and low FRET with the high FRET state corresponding to the non-rotated ribosome, first intermediate corresponding to the initiation specific state, second intermediate state corresponding to the preferred state upon tRNA binding, and low state corresponding to the novel FRET state.

**Figure 4.**
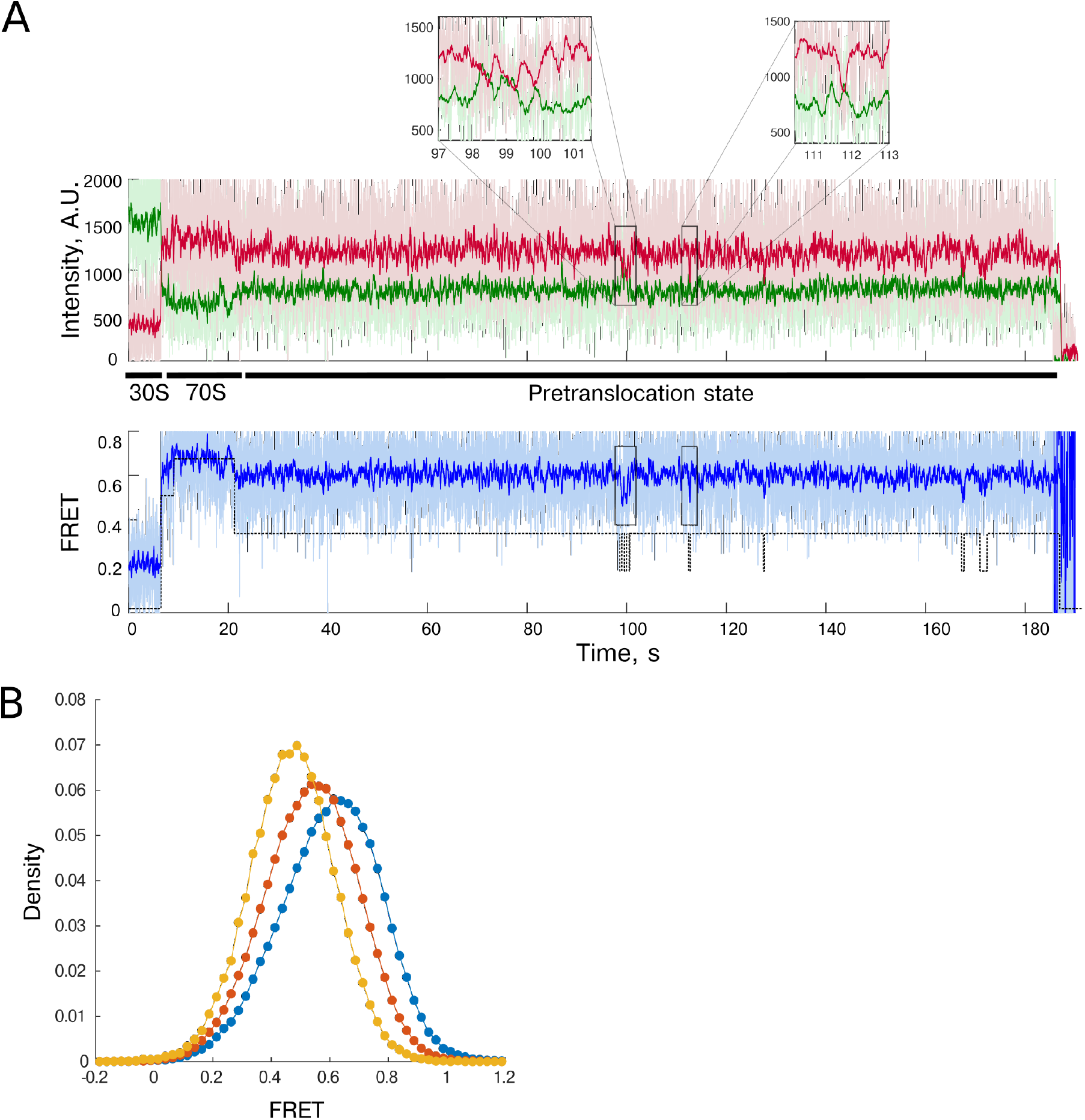
High speed imaging reveals spontaneous rotations in pre-translocation h44-h101 ribosomes. **A.** 50S and TC were delivered to immobilized 30S initiation complexes and imaged at 10 ms resolution. Raw 10 ms fluorescence is shown in light color and 10 frame running average is shown in bold color. Example trace shows partial spontaneous rotations between intermediate and novel low FRET state in pre-translocation ribosomes. **B.** FRET distribution comparison for non-rotated (blue), intermediate (red), and novel low FRET (yellow) states confirms three distinct conformations of the ribosome.

Pre-translocation ribosomes primarily remained in the second intermediate FRET state, with both high and low FRET excursions being short lived, with mean dwell time of 0.9 s (*k_obs_* = 1.1 s^−1^) for the high FRET state and 0.4 s (*k_obs_* = 2.33 s^−1^) for the novel low FRET state (Supplementary Table 2B). The second intermediate state dwell time was best described by double-exponential kinetics. Transitions from the second intermediate to the non-rotated state had a slow phase with a rate of 0.033 s^−1^ and a fast component with a rate of 1.05 s^−1^, while transitions from the second intermediate to low-FRET state had a similar slow phase of *k_obs_*= 0.038 s^−1^, and a fast phase with rate of a 0.20 s^−1^ (Supplementary Table 2C). This complex kinetics are likely due to transient and subtle nature of spontaneous transitions, as many of these transitions possibly remain undetected. However, we could not exclude the possibility that, it reflects a multistep reaction mechanism that underlies reversible spontaneous ribosomal rotations.

To uncover the nature of the novel FRET state we added EF-G to the delivery mix. The EF-G concentration was kept low to prolong the pre-translocation conformation of the ribosome for extended observation. In the presence of EF-G, the ribosomes translocated as evidenced by a reverse transition into a long-lived high-FRET state (Figure 5A, Supplementary Table 3). Importantly, the novel low-FRET state is a better translocation substrate than the second intermediate FRET state. Thirty-six percent of translocation events occurred from the low-FRET state, whereas if both states were equally good substrates the ratio of ribosomes translocated from each state would be proportional to the ratios of low and second intermediate FRET dwell times, i.e. 3%. We post-synchronized all traces to the translocation event, revealing a large density corresponding to the novel state 10-20 ms prior to reverse ribosome rotation (Figure 5E). The prominence of this density, together with its very transient nature, argues that this low state is a ubiquitous translocation substrate and is present even in molecules where it was not directly identified due to the poor signal-to-noise of the high-speed single-molecule trajectories.

**Figure 5.**
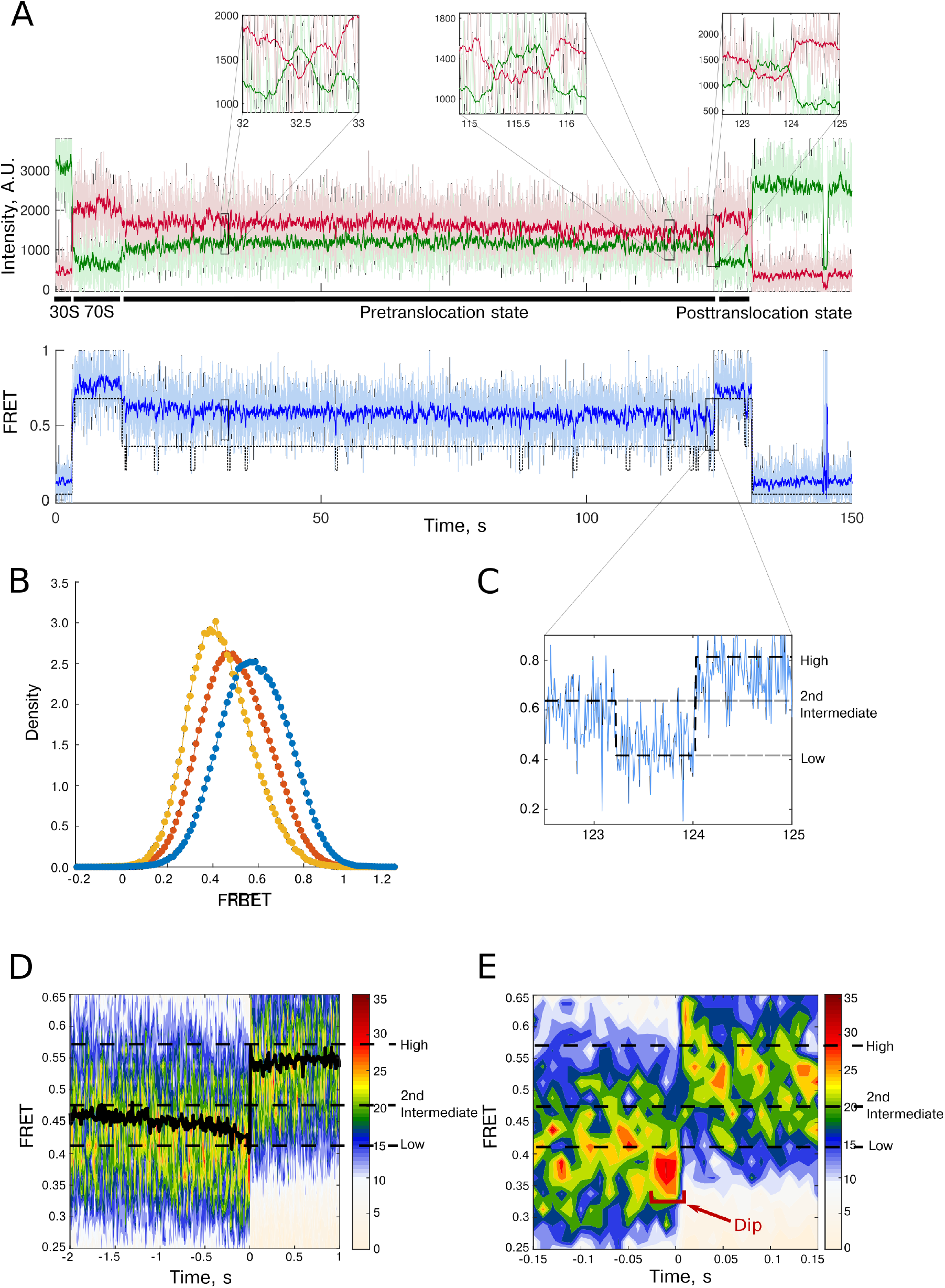
Translocation is preceded by novel low FRET state. Labeled 50S, TC, and EF-G were delivered to immobilized 30S initiation complexes. Translocation was slowed down by low EF-G concentration to extend the pre-translocation state. **A.** Example trace. Pre-translocation ribosomes are predominantly occupying intermediate FRET state, with rapid excursions into novel low FRET state as well as occasional transitions into non-rotated state (Supplementary Figure 7). **B.** FRET distribution comparison of novel state (yellow), intermediate (red) and post translocation non-rotated (blue) ribosomes. **C.** Translocation is preceded by novel low FRET state. In 39% of molecules this state could be directly identified. **D.** All traces were post-synchronized to translocation event. The black line is a median FRET, which decreases right before translocation, further indicating that ribosomes occupy novel conformation. **E.** Zoom into 150 ms prior to translocation shows that ribosomes shift into the novel state for 10-20 ms prior to translocation.

Additionally, we calculated average FRET values at each timepoint and observed the dip in average FRET intensity right before translocation. This dip is analogous to a decrease in S6-L9 FRET that was observed in ensemble kinetics experiments (Belardinelli et al., 2016; Sharma et al., 2016) and was used to argue the presence of spontaneous rotations. Thus, our results fill the gaps between two very different measurement modalities and permit building of a unified model of elongation dynamics.

What is the relationship between FRET states observed in high and low time resolution imaging? Partial spontaneous rotations are transient, which makes them undetectable in low time resolution experiments. The measured FRET values in these experiments are time weighted average of the states detected in high speed experiments. However, due to the transient nature of the spontaneous rotations, their contribution to the measured FRET values is expected to be minor. Therefore, a high FRET state in low time resolution experiments likely corresponds to the high FRET state in high time resolution experiments, which is the predominant state in ribosomes prior to A-site tRNA binding. The low FRET state in the low time resolution experiments corresponds roughly to the second intermediate state in the high time resolution experiments, which is the predominant state in pretranslocation ribosomes.

## Discussion

During translation, bacterial ribosomes adopt at least four different rotational states. Ribosomes join in the first intermediate rotational state. This unique rotational conformation that was not observed during elongation, with rotational state being in between high and second intermediate FRET states observed in pre-translocation ribosomes. GTP hydrolysis by IF2 puts the ribosome into a high FRET state. The high FRET state corresponds to the non-rotated conformation of the ribosome. During elongation, ribosomes undergo exchange between two global states. A-site codon decoding, and translocation separate these two global states. Conformational changes between these global states were used to follow translation over multiple codons. High-speed imaging demonstrated that spontaneous rotations are part of active translation, but are too fast to be detected in conventional singlemolecule translation experiments. Ribosomes experience spontaneous rotations at every step of the translation cycle. Prior to tRNA binding, ribosomes can exchange between high (nonrotated) and second intermediate states. Spontaneous rotations are more common in the pre-translocation portion of the elongation cycle. The conformational exchange in pretranslocation ribosomes occurs between three different states. Ribosomes spend the majority of the time in the second intermediate state, periodically undergoing excursions to the high (non-rotated) and novel FRET states. This novel FRET state is a universal translocation intermediate that precedes reverse intersubunit rotation (Figure 6). What is the nature of the second intermediate and novel low FRET states? It is plausible that the intermediate FRET represents ribosomes that are very rapidly exchanging between high and novel FRET, so the measured FRET is an average of the two states. In this case, the novel FRET state is likely represents fully rotated ribosomal conformation and ribosome fluctuates between rotated and non-rotated conformations during intermediate FRET state. This hypothesis is corroborated by pre-translocation ribosome occupying both of these states, as wells as by transient nature of these rotation intermediates. Congruently, ensemble measurements also predicted rapid exchange between non-rotated and rotated conformations with rates of 40 s^−4^-100 s^−1^(Sharma et al., 2016). This interpretation is also in agreement with spontaneous rotations observed in static experiments (Cornish et al., 2008). The difference in rotation rates in these and our experiments are likely due to experimental conditions and varying concentrations of polyamines and magnesium as was previously proposed (Sharma et al., 2016).

**Figure 6.**
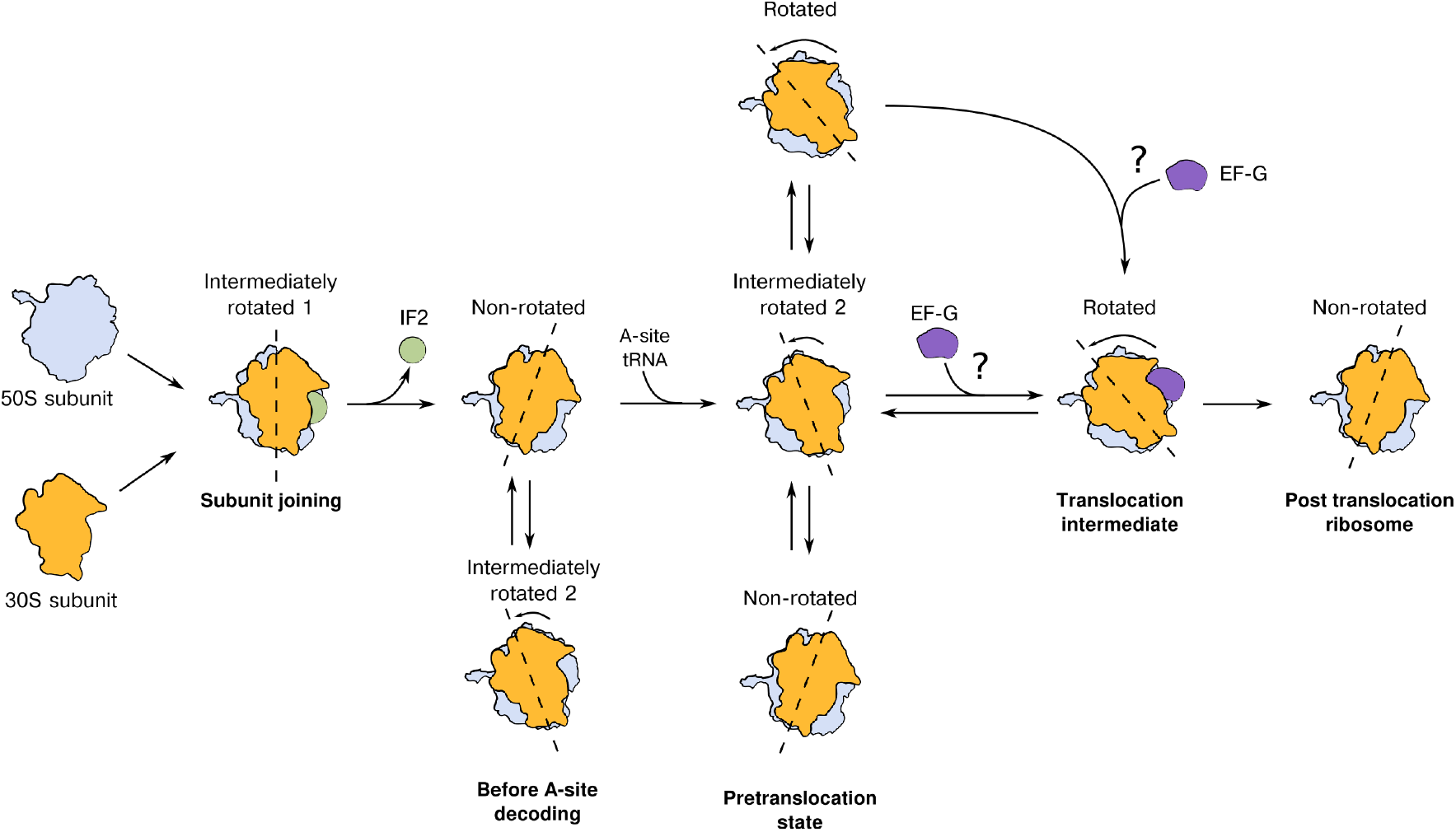
Model of ribosome movements during translation elongation. Ribosomes initially join in a partially rotated state. Rapid GTP hydrolysis by IF2 triggers the transition into a non-rotated state. While awaiting A-site tRNA decoding, the ribosome predominantly occupies a non-rotated conformation with occasional brief excursions into a second intermediate rotational state. After A-site tRNA binding and peptidyl transfer, the ribosome predominantly occupies this second intermediate rotational state. Pre-translocation ribosomes can undergo spontaneous transitions into a non-rotated, as well as into a novel rotational state, that presumably represent the fully rotated ribosome. This state is also a translocation intermediate, as the ribosome adopts a fully rotated conformation for 10-20 ms prior to reverse intersubunit rotation catalyzed by EF-G. Translocation results in the non-rotated ribosomes.

The ribosome occupies a novel low-FRET state prior to reverse rotation. This is possibly a cause of the FRET decrease prior to translocation observed in ensemble experiments. Similarly, it was interpreted as ribosomes that fluctuate between rotated and non-rotated states are being biased into the rotated state by EF-G binding.

Alternatively, the three states observed during elongation could be individual rotational states with the novel state representing either fully rotated or hyper-rotated ribosomes. It was shown that mRNA structure and SecM stall sequence might induce hyper-rotated conformation (Klimova et al., 2019; Qin et al., 2014). Our results suggest that these events occur via a rotational state that represents a stalled translocation intermediate.

If ribosome occupies three distinct states, it spontaneously transitions between fully and partially rotated states in the pre-translocation state. In this model, during translocation EF-G promotes partial intersubunit rotation into rotated or hyper-rotated state before catalyzing full reverse rotation. Additional improvements in the time resolution are needed to distinguish between these possibilities.

### Future directions

In vivo translation occurs rapidly occurring at rates of ~ 10 amino acids per second. These measurements approach the resolution needed to observe translation at in vivo speeds. High speed imaging could be further applied to study all phases of translation. This includes regular translation, as well as regulatory and decoding events that undergo via unstable intermediates that are governed by kinetic partitioning.

## Acknowledgments

This work was supported by Auburn University PAIR grant to A.P and NIH grant GM51266 to JDP. We thank H. Noller and L. Lancaster for providing labeled ribosomes, and many useful discussions. We thank Brielle Sorkin for helping with manuscript preparation.

## Contributions

AP and JDP conceived the study. AP conducted low time-resolution experiments. TH performed low and high-temporal resolution experiments. AP and TH analyzed the data. JC and ArPr provided critical reagents and insight into experimental design and planning. AP and TH wrote the manuscript with help and editorial input from all other authors. AP and JDP supervised the research.

---

**Supplementary Figure 1.**
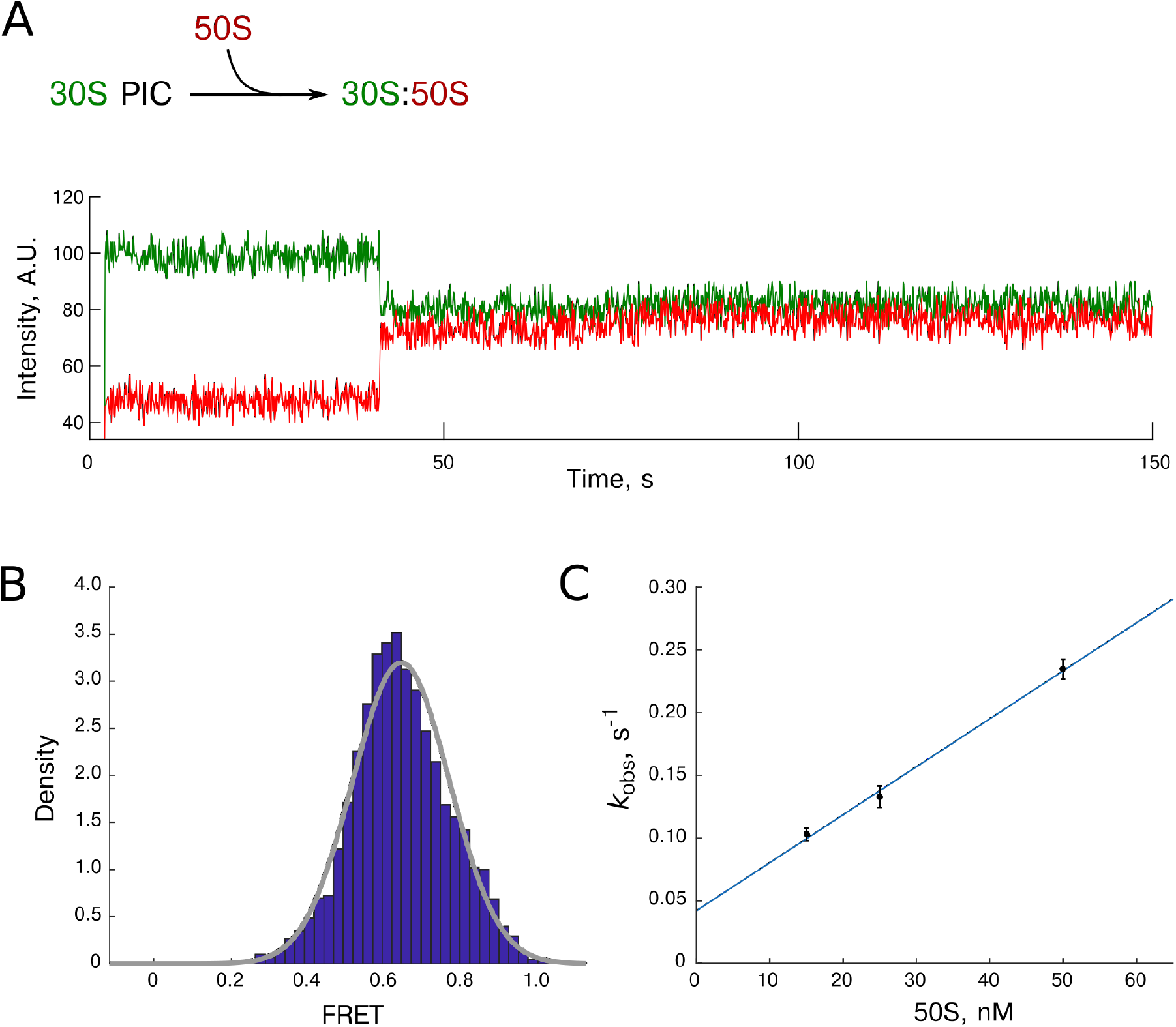
Following 70S formation in real time using S6-L9 FRET. 30S initiation complexes comprised of 30S-Cy3 ribosomal subunits (labeled at ribosomal protein S6), initiator tRNA, and IF2 were assembled over biotinylated (FK)_6_ mRNA. Complexes were immobilized on the surface. 50S-Cy5 ribosomal subunits (labeled at ribosomal protein L9) were delivered briefly after the start of imaging. **A.** Example trace of 50S-Cy5 delivery to 30S PICs (30S-Cy3:fMet-tRNA^fMet^:IF2:mRNA-biotin). It begins with 30S-Cy3 fluorescence from immobilized 30S PICs. 50S subunit binds at ~ 40 s, which is indicated by the FRET appearance. FRET efficiency is stable until the end of acquisition. **B.** FRET efficiency histogram is single modal. Cryo-EM and X-ray structure showed 70S ribosomes with fMet-tRNA^fMet^ in the P-site occupying non-rotated conformation. Thus we attribute this state to the non-rotated ribosomes. The gray line represents a single Gaussian fit. **C.** The subunit-joining rate was concentration-dependent. N = 89, 1617 and 378 correspondingly. Error bars are errors of the fit. The rate constant was determined as the slope of the concentration curve *k*_on_ = 3.8±0.4 μM^−1^s^−1^. The y-axis intercept is 30S-Cy3 photobleaching rate of 0.042 s^−1^, which closely corresponds to the directly measured value of 0.057±0.004 s^−1^, n = 222.

**Supplementary Figure 2.**
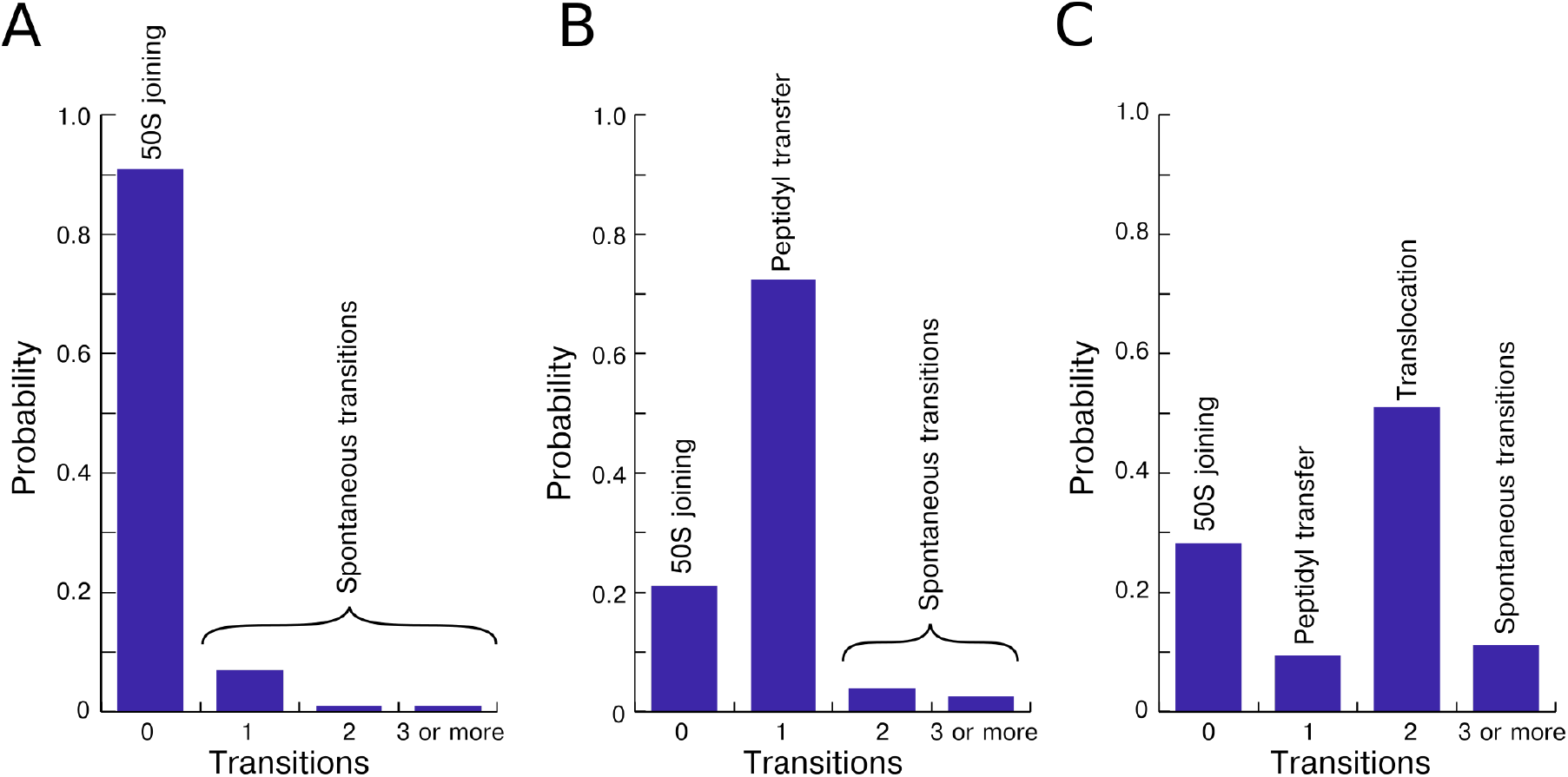
Spontaneous inter-subunit rotations in actively translating S6-L9 ribosomes. **A.** Number of rotations observed in 50S subunit joining experiments. The majority of ribosomes (91%) did not show any change in FRET values. Seven percent spontaneously switched from high FRET to the low FRET once. One percent of the molecules underwent a full spontaneous rotation cycle, first moving into the rotated state and then back into the non-rotated state. Additionally, 1% of ribosomes demonstrated multiple inter-subunit movements. **B.** Number of ribosome rotations in 50S subunit and ternary complex delivery experiments. 21 percent of the molecules underwent a subunit joining and had a stable FRET. There are two kinds of molecules in this category. First are molecules where the dyes photobleached before the first elongation codon was read. These are active molecules where ribosomal rotations could not be observed. Second are inactive molecules that stack on the 70S initiation complex stage. 72 percent of the molecules transitioned from high-to-low FRET, consistent with the reading of the first elongation (Phe) codon. Four percent of the ribosomes experienced a single spontaneous transition back into the non-rotated state. 3 percent of the ribosomes underwent multiple spontaneous rotations. **C.** Number of intersubunit rotations in presence of Phe-tRNA^Phe^ and EF-G. Most ribosomes (51%) showed two rotations consistent with A-site tRNA binding and translocation. The first rotation occurred upon tRNA binding from the non-rotated state to the rotated state. The second rotation occurred from the rotated state to the non-rotated state upon translocation. Additionally, 11 percent of the ribosomes demonstrated extra inter subunit rotations. Ribosomes with 0 or 1 transitions are a combination of ribosomes that either photobleached before they underwent peptidyl transfer (0 transitions) or translocation (1 transition) and inactive ribosomes.

**Supplementary Figure 3.**
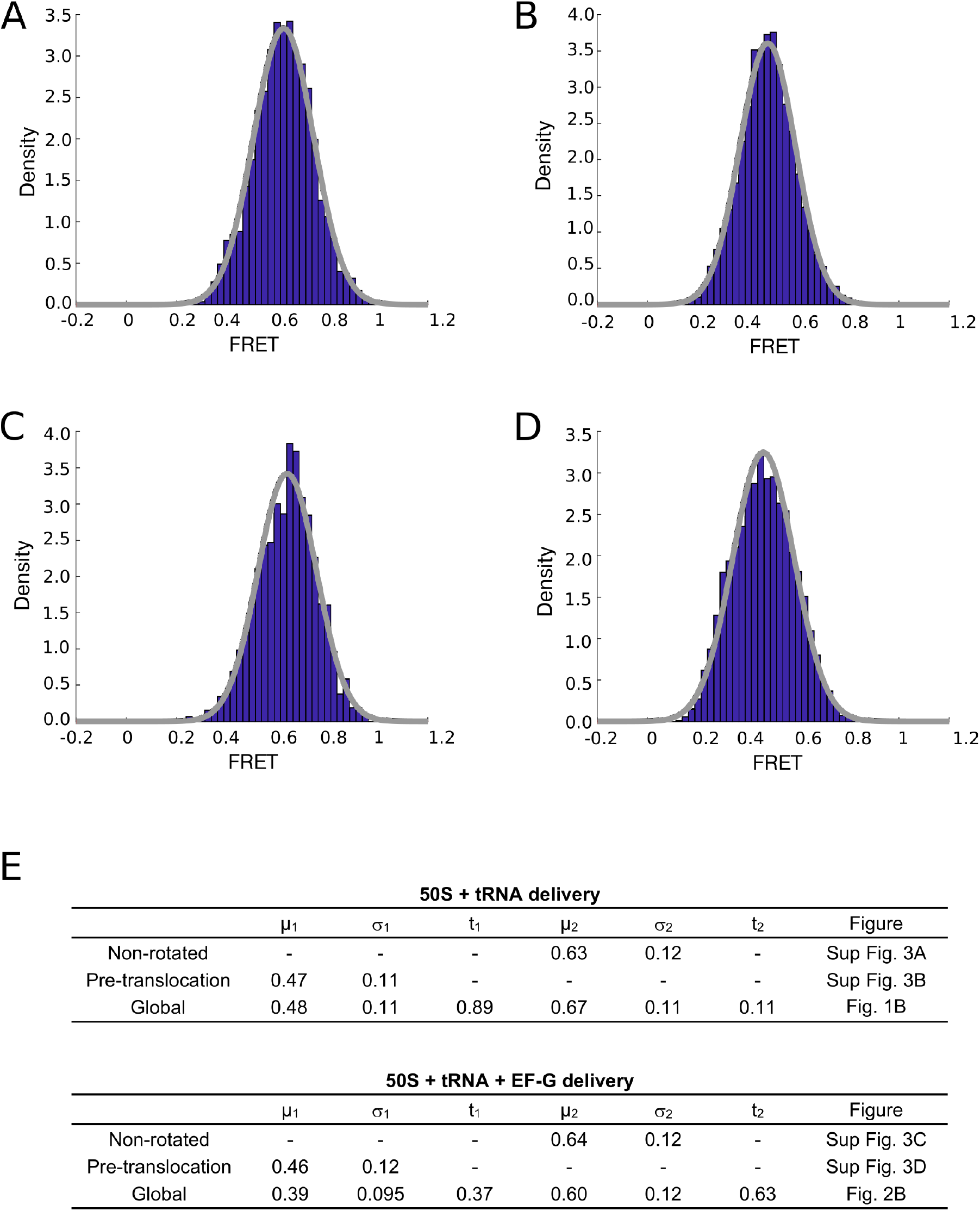
FRET intensity distributions and Gaussian fit results. **A and B** show separate FRET intensity distributions for each individual state in 50S subunits and tRNA delivery experiment. **A.** FRET intensity histogram for high FRET (nonrotated ribosomes) **B.** FRET distribution for pre-translocation ribosomes. **C and D** show FRET intensity distributions for each state in 50S, tRNA and EF-G delivery experiment. FRET distribution for high FRET (non-rotated ribosomes). This state is comprised of ribosomes after subunit joining awaiting A-site decoding and post-translocation ribosomes. **D.** FRET distribution for pre-translocation ribosomes. **E.** Results of double Gaussian fits of global FRET distributions and single Gaussian fits of distributions representing individual functional states. Tables show μ (mean), σ (standard deviation) and t (probability) for each Gaussian.

**Supplementary Figure 4.**
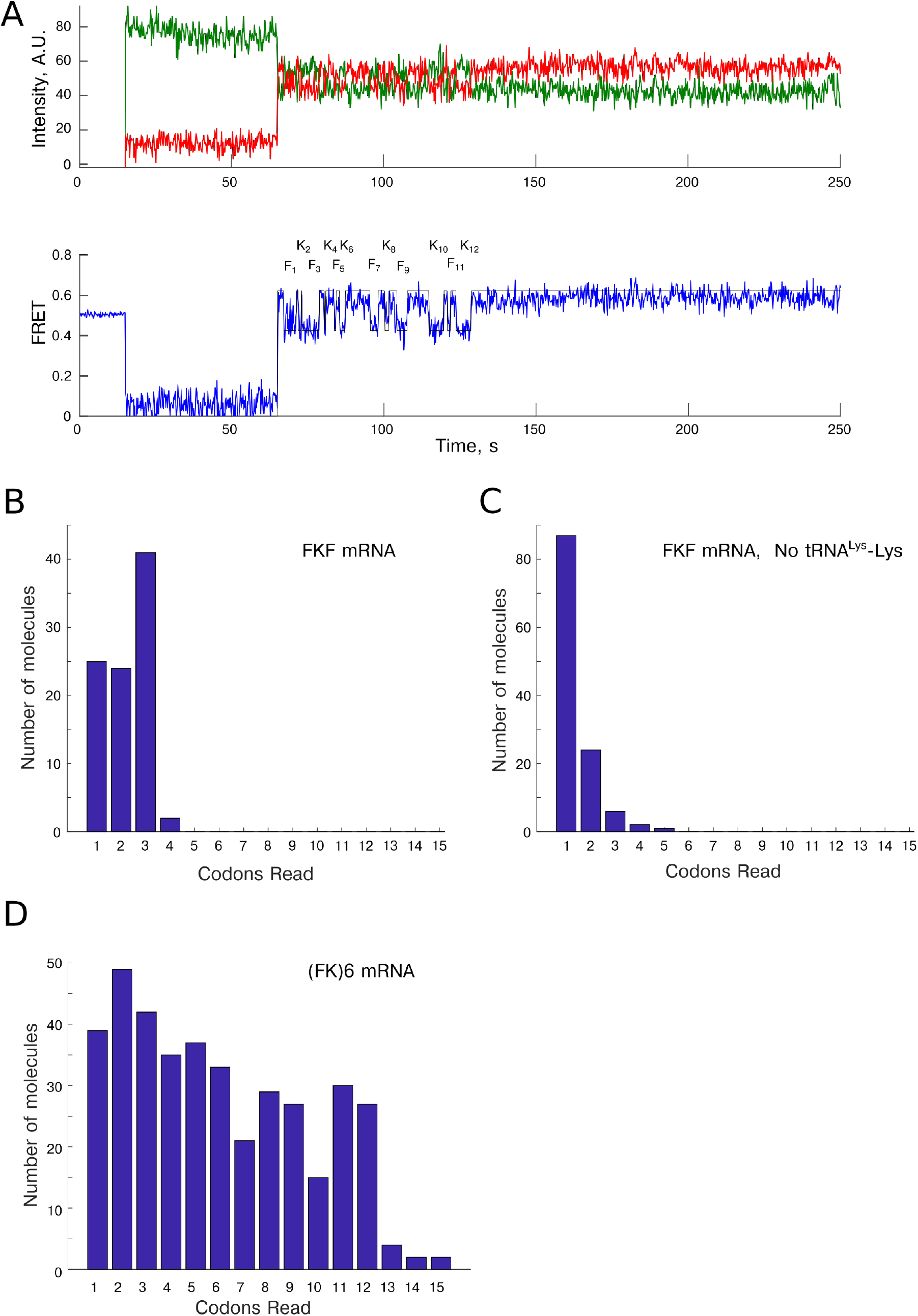
Number of codons read on FKF and (FK)6 mRNA. The number of codons read was determined as a number of full cycles of ribosome rotations. Number of observed ribosomal rotations strongly correlates with number of translated codons. **A.** Example trace of translation of (FK)6 mRNA. Subunit joining occurs at around 50 s mark. It is followed by the rapid cycling from rotated to non-rotated states and back, for a total of twelve cycles. The trace idealization used for data analysis is shown by a hairline in the bottom FRET panel. The codon identity is indicated. **B.** Number of codons read on FKF mRNA. Accumulation of the ribosomes on the last, third codon, is apparent with only 2% of the ribosomes experiencing more than 3 rotation cycles. N = 92. **C.** Number of codons read on FKF mRNA in the absence of Lys-tRNA^Lys^. The majority (79%) of ribosomes underwent a single rotational cycle, n=110. **D.** Number of translated codons on (FK)6 mRNA. Translation of up to 12 codons was observed. Photobleaching severely limits the number of observed codons. Five percent of the ribosomes underwent more than 12 rotational cycles. They represent a sum of a) freely fluctuating ribosomes b) error in rotation assignment c) ribosomes that read through the stop codon. N = 412.

**Supplementary Figure 5.**
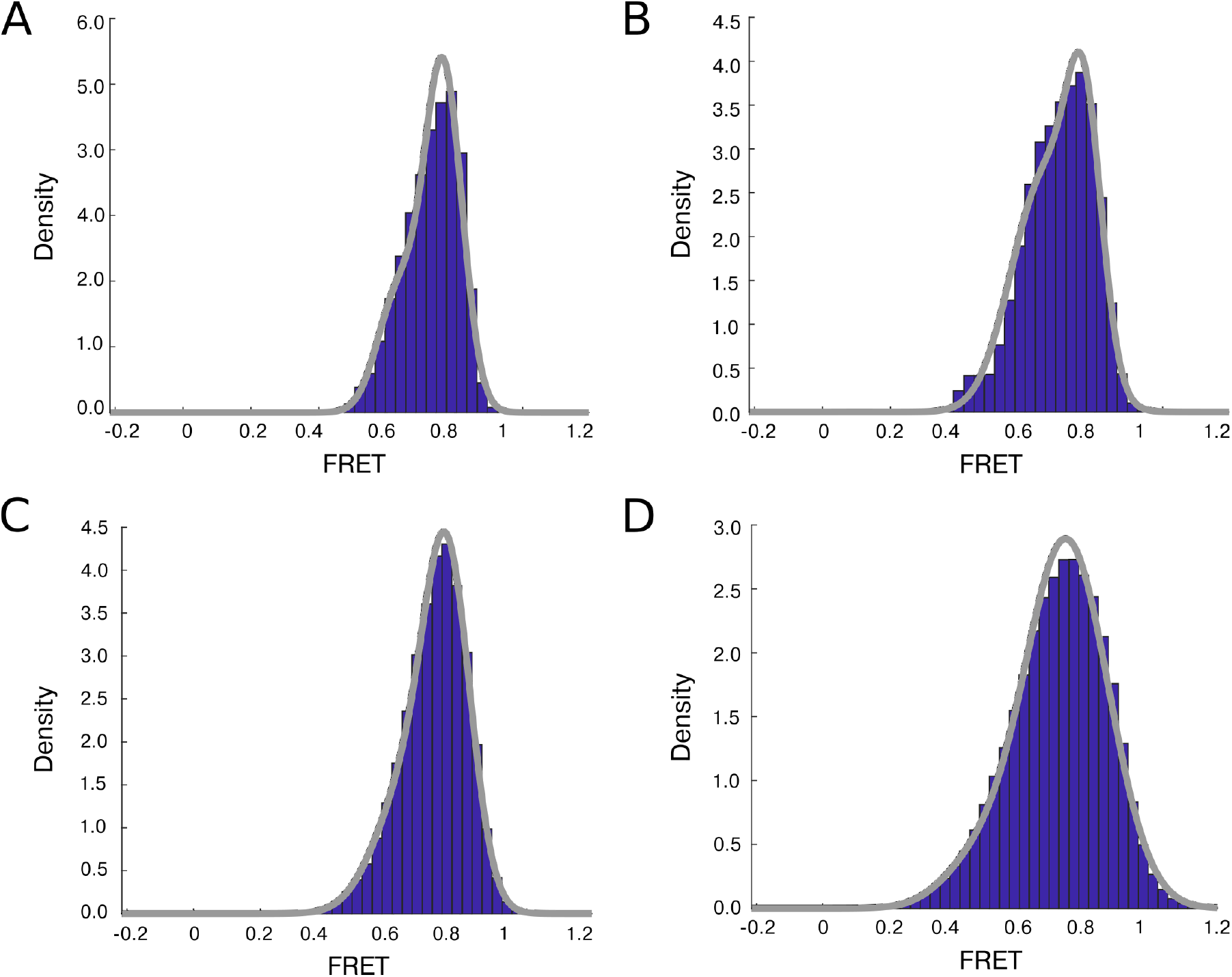
Characterization of high-speed single-molecule microscope. DNA oligonucleotide carrying 5’ Cy3 and 3’ biotin was annealed to the DNA oligonucleotide carrying 5’ Cy5. The resulting duplex was immobilized and imaged at different experimental conditions. **A.** FRET intensity histogram at 200 ms exposure. These are conditions routinely used in single molecule experiments. **B.** FRET intensity distributions at 50 ms exposure time. **C.** FRET intensity distributions at 25 ms exposure time. There is a negligible increase in signal-to-noise up to 25 ms resolution. **D.** FRET intensity histogram at 10 ms exposure used in this work.

**Supplementary Figure 6.**
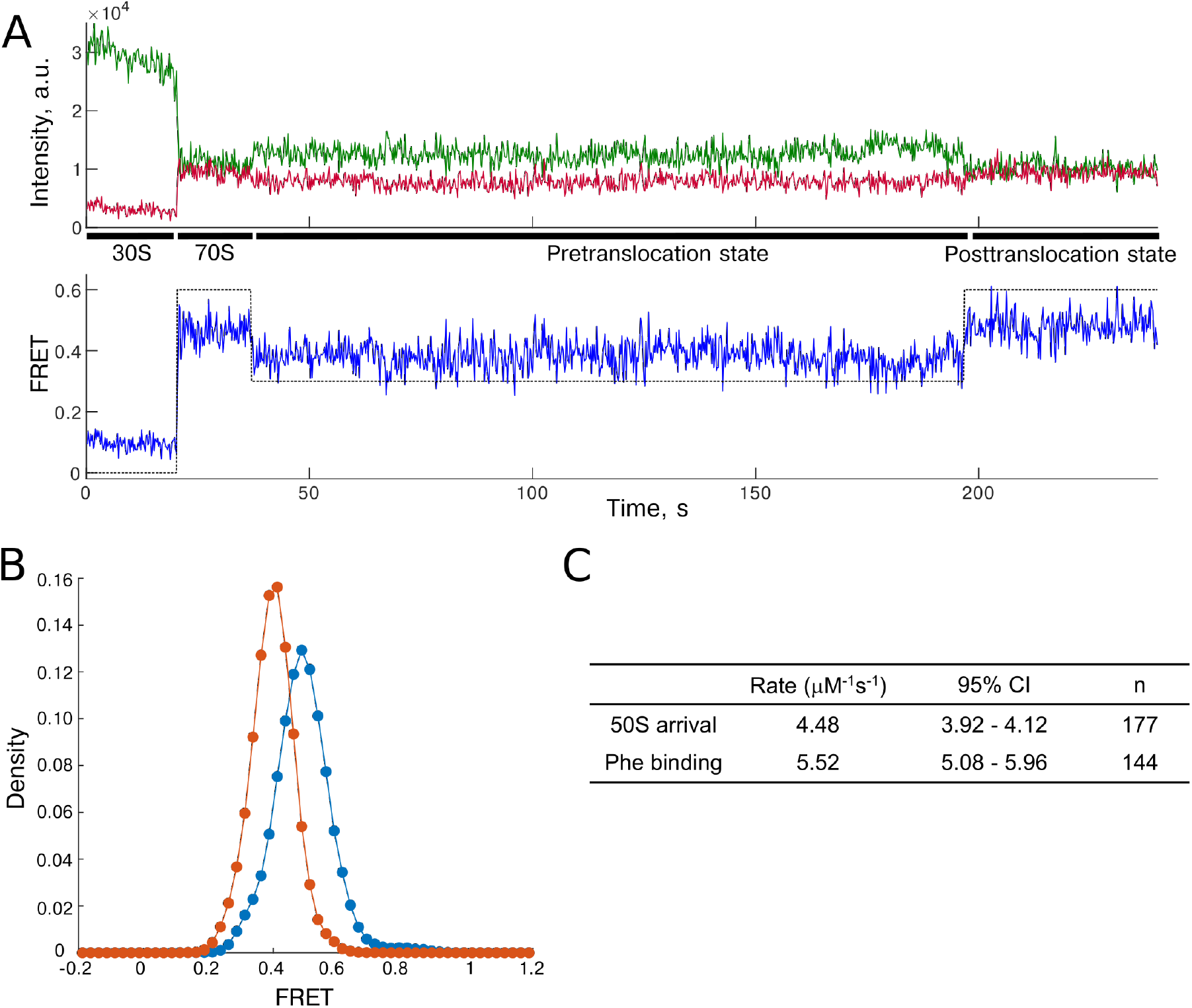
Translation at low-speed conditions with h44-h101 labeled ribosomes. 50S subunits, TC, and EF-G were delivered to immobilized 30S initiation complexes and translation was imaged at 200 ms resolution. **A.** Example trace. Ribosomes alternate between two global states upon tRNA binding and translocation. **B.** FRET intensity distributions for post-translocation (blue, FRET intensity of 0.50) and pre-translocation ribosomes (red, FRET intensity of 0.40). **C.** Observed rates of 50S subunit binding and A-site tRNA arrival, 95% CI is 95% confidence interval.

**Supplementary Figure 7.**
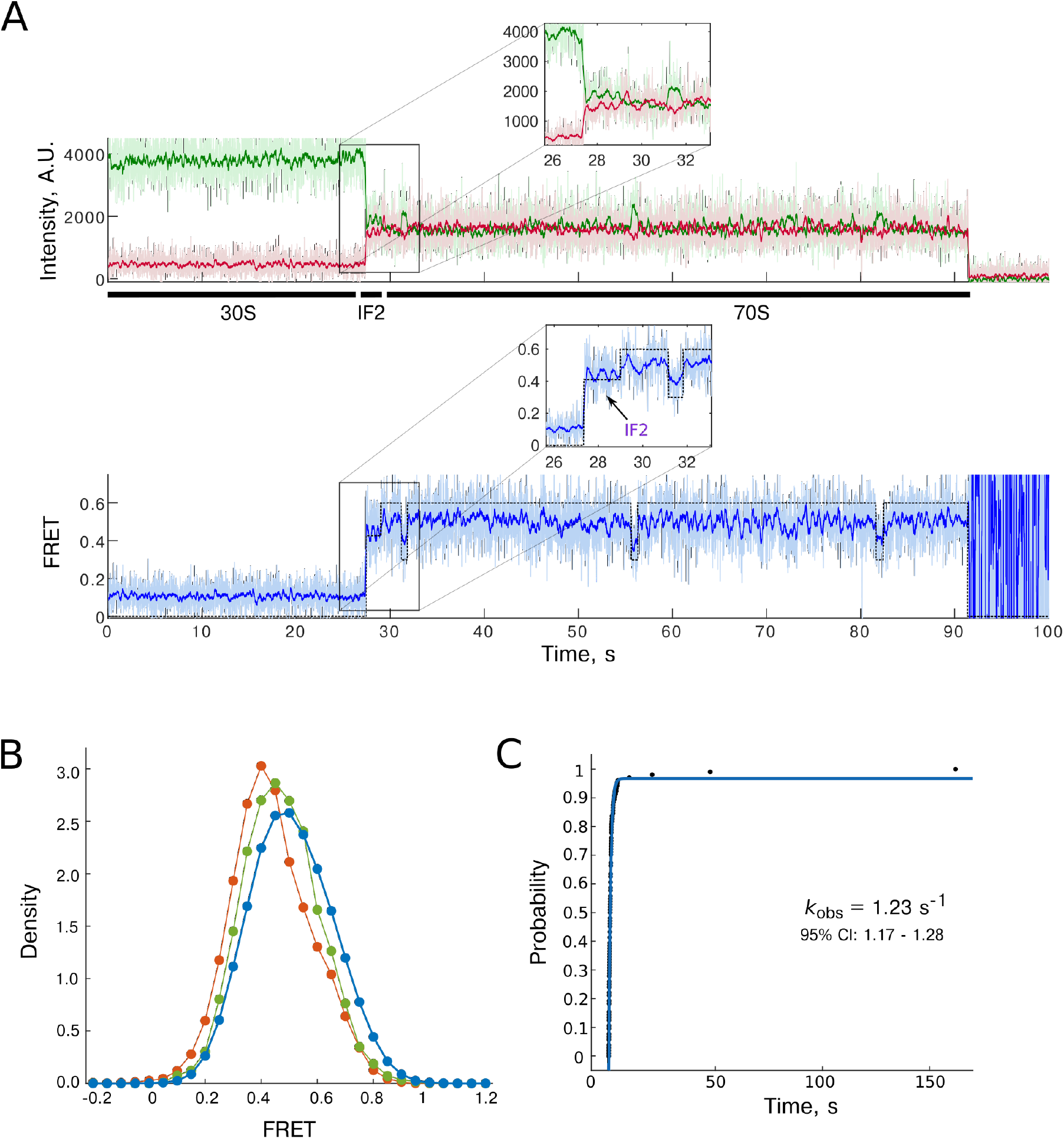
Following initiation with high temporal resolution. 50S subunits were delivered to surface immobilized 30S PIC and imaged at 10 ms resolution. **A.** Example trace. Raw 10 ms fluorescence is shown in light color and 10 frame running average is shown in bold color. Subunits joined in the intermediate rotation state. GTP hydrolysis by IF2 finalizes initiation and shifts ribosome into the non-rotated state. Ribosome undergoes spontaneous rotations (32 and 56 second marks) while it awaits A-site tRNA. **B.** FRET intensity distribution comparison: state before GTP hydrolysis by IF2 (green; FRET intensity of 0.47), spontaneous transitions in 70S ribosomes, same as the second intermediate state (red; mean of 0.44), and a non-rotated state (blue, with mean equal to 0.51). Subunits join in the intermediate rotated state that is different from both nonrotated and spontaneously rotated (same as second intermediate state, see Figure 5) ribosomes. **C.** Rotation state dwell times. While occasional long-lived spontaneous rotations have been observed, the majority of rotations are transient.

**Supplementary Figure 8.**
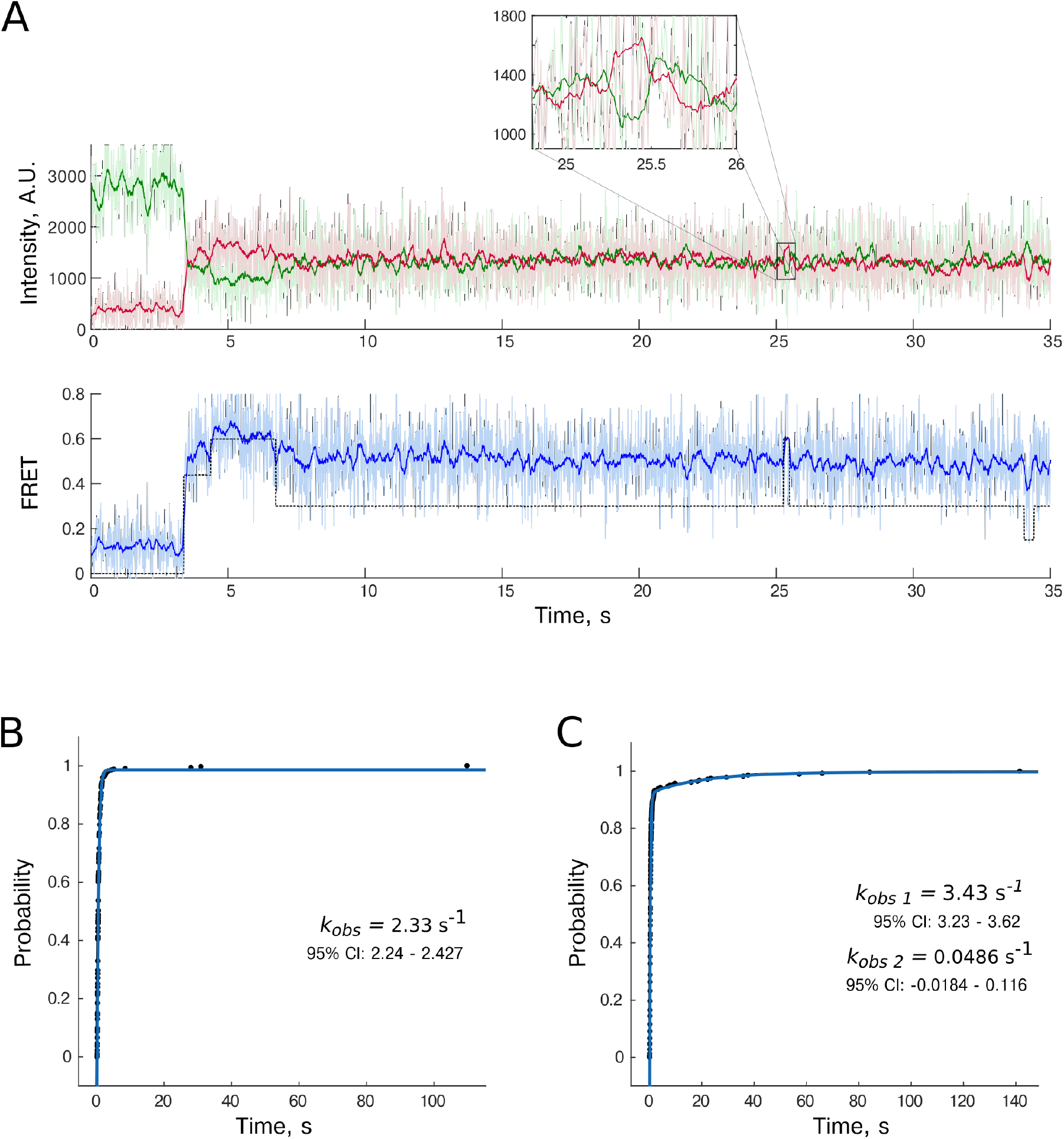
Partial rotations from 2^nd^ intermediate to high FRET state (non-rotated conformation) in pre-translocation ribosomes. Further analysis of the experiment described in Figure 4 revealed that ribosomes can undergo transient excursions into a non-rotated state. **A.** Example trace. Spontaneous transition occurs at the 25 s mark. Rotations occurred spontaneously and did not require EF-G. **B.** Kinetics of spontaneous transitions from 2^nd^ intermediate state into high FRET state in pre-translocation ribosomes in 50S subunits and TC delivery experiment. **C.** Kinetics of spontaneous transitions from 2^nd^ intermediate state into high FRET state in pretranslocation ribosomes in 50S subunits, TC, and EF-G delivery experiment.

**Supplementary Table 1.**
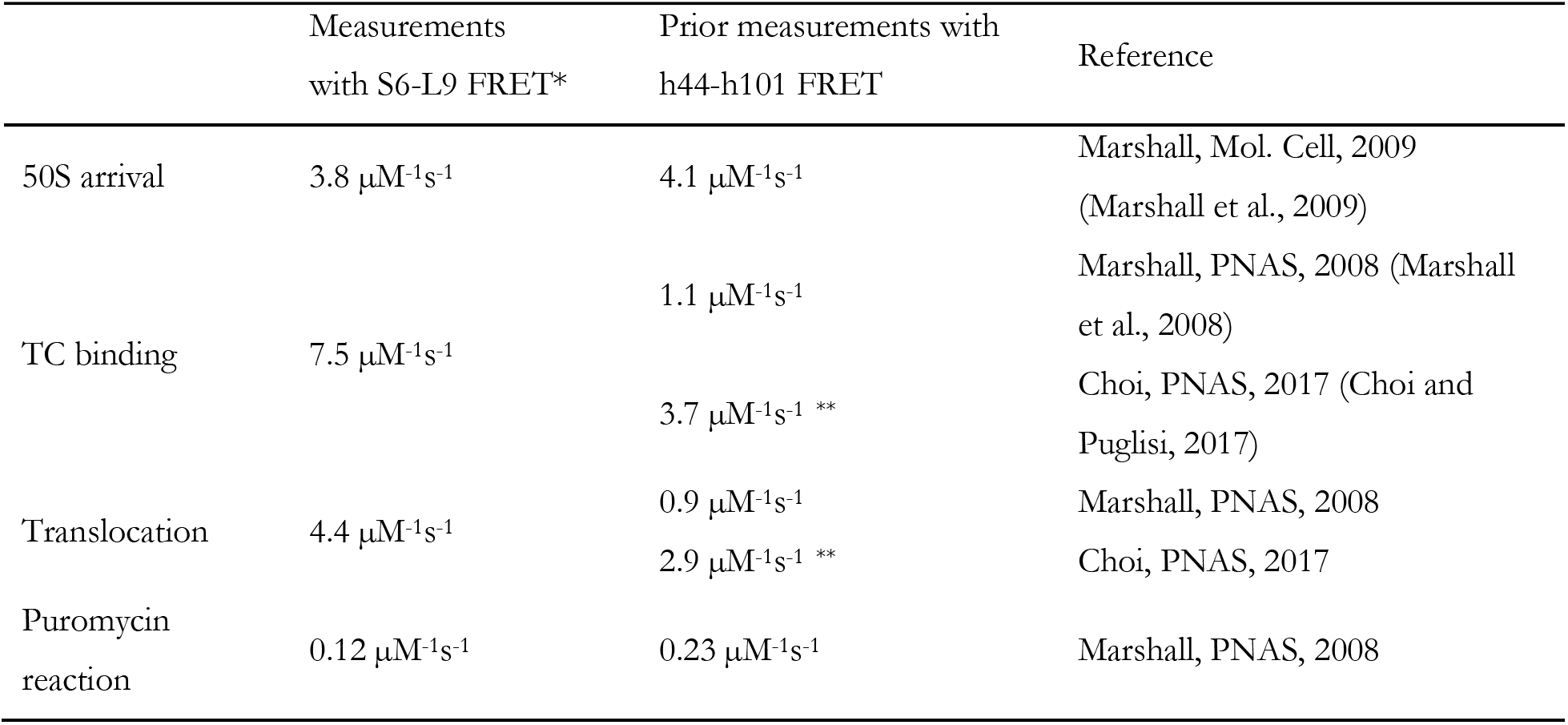
Comparison of translation kinetics between h44-h101 and S6-L9 ribosomes. *The kinetic parameters were calculated as a slope of concentration vs reaction rate charts. ** Kinetic parameters were calculated from data presented in (Choi and Puglisi, 2017). Average TC binding time of 13.6 s at 20 nM tRNA yielded a 3.7 μM^−1^s^−1^ reaction rate constant for A-site tRNA decoding. Average dwell time in the pre-translocation state of 8.5 s at 40 nM EF-G is equivalent to the 2.9 μM^−1^s^−1^ reaction rate constant.

|  | Measurements<br>with S6-L9 FRET* | Prior measurements with<br>h44-h101 FRET | Reference |
| --- | --- | --- | --- |
| 50S arrival | 3.8 $\mu\text{M}^{-1}\text{s}^{-1}$ | 4.1 $\mu\text{M}^{-1}\text{s}^{-1}$ | Marshall, Mol. Cell, 2009<br>(Marshall et al., 2009) |
| TC binding | 7.5 $\mu\text{M}^{-1}\text{s}^{-1}$ | 1.1 $\mu\text{M}^{-1}\text{s}^{-1}$ | Marshall, PNAS, 2008 (Marshall<br>et al., 2008) |
| | | 3.7 $\mu\text{M}^{-1}\text{s}^{-1}$ ** | Choi, PNAS, 2017 (Choi and<br>Puglisi, 2017) |
| Translocation | 4.4 $\mu\text{M}^{-1}\text{s}^{-1}$ | 0.9 $\mu\text{M}^{-1}\text{s}^{-1}$ | Marshall, PNAS, 2008 |
| | | 2.9 $\mu\text{M}^{-1}\text{s}^{-1}$ ** | Choi, PNAS, 2017 |
| Puromycin<br>reaction | 0.12 $\mu\text{M}^{-1}\text{s}^{-1}$ | 0.23 $\mu\text{M}^{-1}\text{s}^{-1}$ | Marshall, PNAS, 2008 |

**Supplementary Table 2.**
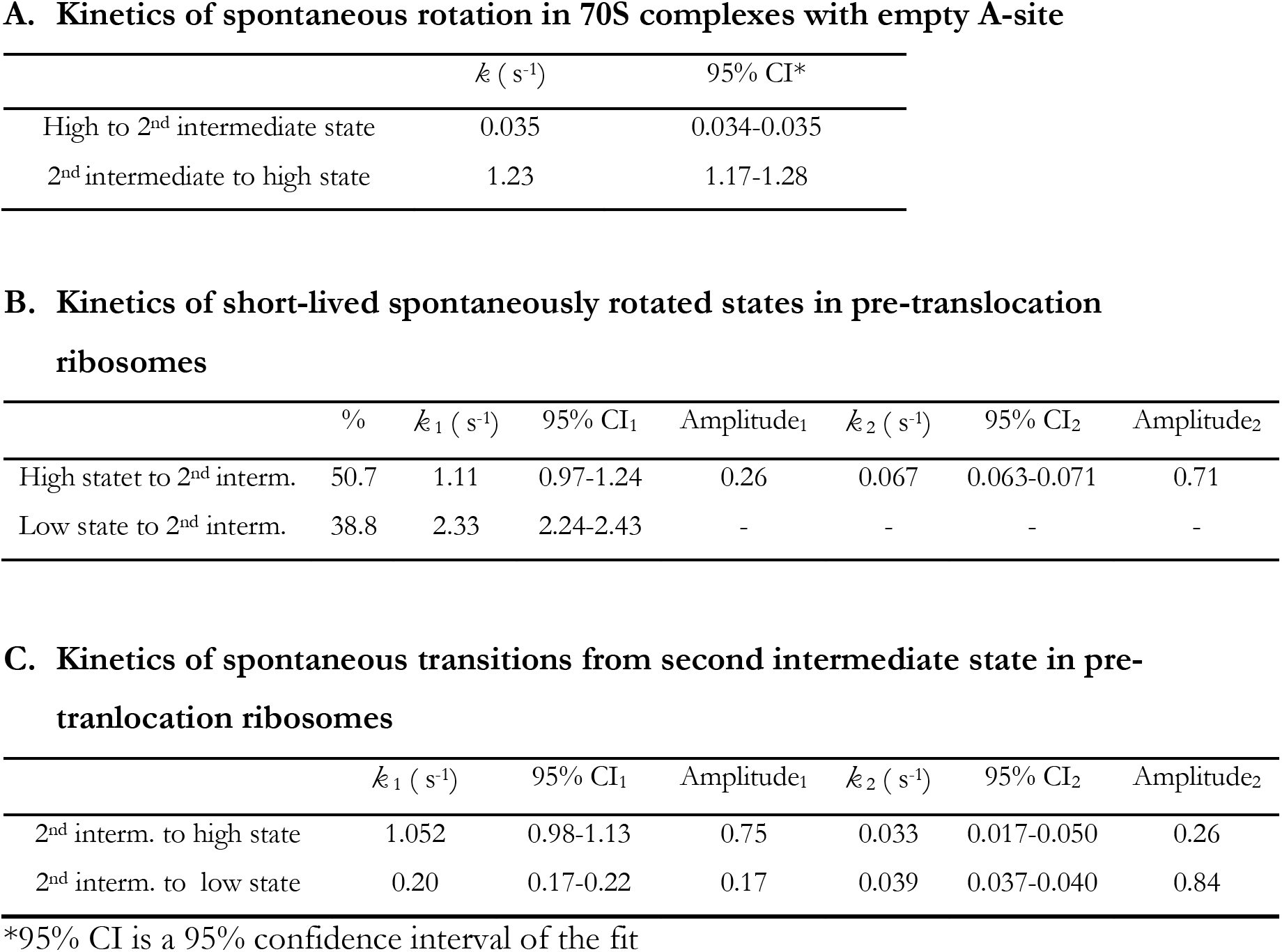
**A.** Kinetics of spontaneous rotation in 70S complexes with empty A-sites. NR is a nonrotated high FRET state. 2^nd^ intermediate state is a state in which pre-translocation ribosomes spend the majority of time. N=208. **B.** Transition rates from short lived high and low FRET states into the second intermediate state in pre-translocation ribosomes. Shows kinetics of spontaneous rotations from the intermediate state to the novel low and nonrotated states. N=209. **C.** Reaction rates that govern second intermediate state dwell times. Shows kinetics of spontaneous rotation from the intermediate state to the low or high FRET states.

A. Kinetics of spontaneous rotation in 70S complexes with empty A-site
| | $k$ (s<sup>-1</sup>) | 95% CI* |
| --- | --- | --- |
| High to 2<sup>nd</sup> intermediate state | 0.035 | 0.034-0.035 |
| 2<sup>nd</sup> intermediate to high state | 1.23 | 1.17-1.28 |

B. Kinetics of short-lived spontaneously rotated states in pre-translocation ribosomes
| | % | $k_1$ (s<sup>-1</sup>) | 95% CI<sub>1</sub> | Amplitude<sub>1</sub> | $k_2$ (s<sup>-1</sup>) | 95% CI<sub>2</sub> | Amplitude<sub>2</sub> |
| --- | --- | --- | --- | --- | --- | --- | --- |
| High statet to 2<sup>nd</sup> interm. | 50.7 | 1.11 | 0.97-1.24 | 0.26 | 0.067 | 0.063-0.071 | 0.71 |
| Low state to 2<sup>nd</sup> interm. | 38.8 | 2.33 | 2.24-2.43 | - | - | - | - |

Supplementary Table 2.
| | $k_1$ (s<sup>-1</sup>) | 95% CI<sub>1</sub> | Amplitude<sub>1</sub> | $k_2$ (s<sup>-1</sup>) | 95% CI<sub>2</sub> | Amplitude<sub>2</sub> |
| --- | --- | --- | --- | --- | --- | --- |
| 2<sup>nd</sup> interm. to high state | 1.052 | 0.98-1.13 | 0.75 | 0.033 | 0.017-0.050 | 0.26 |
| 2<sup>nd</sup> interm. to low state | 0.20 | 0.17-0.22 | 0.17 | 0.039 | 0.037-0.040 | 0.84 |

\*95% CI is a 95% confidence interval of the fit

**Supplementary Table 3.** FRET values in high-speed translation experiments. Due to high signal to noise, absolute FRET values showed a significant degree of variability, with absolute values varying as much as 0.1. Thus, we used a combination of absolute FRET values with event sequences and to identify corresponding translational events and translation intermediates. The FRET intensity that is characterizes characteristic to each state was determined using two approaches: First, FRET efficiencies were obtained as a mean of a single Gaussian fit using an expectation-minimization algorithm. These values are reported in “Gaussian” rows. Notably, FRET intensity distributions are asymmetric. Consequently, means of Gaussian fits were not coinciding with the modes of the distributions. Thus, a second approach was used: modes of FRET intensity distributions were determined as modes of FRET intensity histograms built with a 0.0125 step. These values are reported in the “Mode” rows and used in the text.

| Experiment | Method | High FRET<br>(Non-rotated) | 1<sup>st</sup> intermed.<br>(Initiation<br>specific) | 2<sup>nd</sup> intermed.<br>(Major pre-translocation) | Low FRET<br>(Novel state) |
| --- | --- | --- | --- | --- | --- |
| 50S delivery to | Gaussian | 0.51 | 0.47 | 0.44 | - |
| 30S PICs | Mode | 0.50 | 0.45 | 0.40 | - |
| 50S and tRNA | Gaussian | 0.60 | - | 0.54 | 0.47 |
| delivery to 30S PICs | Mode | 0.63 | - | 0.53 | 0.48 |
| 50S, tRNA and EF-G | Gaussian | 0.58 | - | 0.50 | 0.43 |
| delivery to 30S PIC | Mode | 0.59 | - | 0.49 | 0.42 |

## Materials and methods

### Reagents

30S ribosomal subunits labeled at D41C of RPS6 and 50S subunits labeled at N11C of RPL9 were prepared as previously described (Ermolenko et al., 2007). Small and large ribosomal subunits labeled at rRNA extensions (h44 and h101 correspondingly) were purified and labeled with fluorescent oligonucleotides as previously described. Briefly, the apical loops of h44 of 16S rRNA and h101 of 23S rRNA were replaced with labile hairpins (Dorywalska et al., 2005; Marshall et al., 2008). The short oligonucleotides derivatized with fluorescent dyes Cy3-GAGGCCGAGAAGTG and Cy5-GGGAGATCAGGATA were annealed to the hairpins, resulting in site-specifically labeled ribosomal subunits. Translation factors, mRNA and tRNAs were obtained as before (Aitken and Puglisi, 2010; Blanchard et al., 2004). Both FKF and (FK)6 mRNAs have a 5’ UTR and Shine-Dalgarno sequence derived from gene 32 of the T4 phage. In (FK)6 mRNA coding sequence consists of initiation AUG codon and six repeats of Phe and Lys codons. The open reading frame is terminated with UAA stop codon, which is followed by twelve spacer uridine bases. The full sequence of (FK)6 mRNA is 5’-CAACCUAAAACUUACACACGCCCCGGUAAGGAAAUAAAAAUGUUCAAAUUCAAAUUCAAAUUCAAAUUCAAAUUCAAAUAAUUUUUUUUUUUU-3’. All mRNAs carry 5’ biotin that allows surface immobilization of ribosomal complexes. The FKF mRNA is the same except the six FK repeats are replaced with FKF sequence.

### Real-time tracking of ribosomal conformation in actively translating ribosomes

The single-molecule fluorescence imaging was performed using a home-built prism-based TIRF system (Aitken et al., 2008) and modified commercial ZMW RSII sequencer from Pacific Bioscience (Chen et al., 2014). All reactions were done in polymix buffer containing 50 mM Tris-Acetic acid, pH = 7.5, 100 mM KCl, 5 mM NH_4_OAc, 5 mM Ca(OAc)_2_, 5 mM putrescine, 1 mM spermidine and 0.1 mM EDTA. 30S initiation complexes were assembled over mRNA in a stepwise manner. First, 0.5 μM 30S ribosomes were co-incubated with 0.5 μM ribosomal protein S1 at 37°C for 5 min. Next, the 30S PICs were formed by incubating 0.25 μM 30S+S1, 1 μM mRNA, 1 μM IF2 (1.2 μM in 10 ms experiments), 1 μM fMet-tRNA^fMet^ and 1 mM GTP, in polymix buffer. Both TIRF and ZMW microscopy were done as before (Chen et al., 2014; Marshall et al., 2009). Briefly, the TIRF microscopy was done using in house made microscope chambers of ~ 10 μl volume. 30S PICs complexes were immobilized on the surface of the neutravidin-covered quartz slides at 30 pM for 5 min. The unreacted 30S subunits, complexes lacking mRNA and unbound complexes were washed away with 200 μl of the reaction buffer supplemented with 1 μM IF2, 2.5 mM TSQ (Triplet State Quencher, Pacific Biosciences), 2.5 mM protocatechuic acid, and 1x protocatechuate-3,4-dioxygenase from Pacific Biosciences. The 40 μl delivery mix containing 50S subunits, ternary complex of EF-Tu:GTP with elongator tRNAs and EF-G was delivered simultaneously with data acquisition. The delivery mixture fully replaced the reaction buffer in the chamber resulting in ribosome and factor concentrations as indicated in text. The dyes were excited with a 532 nm laser at approximately 400W/cm^2^ intensity. About 1000 μm^2^ area was imaged and the Cy3 and Cy5 fluorescence was recorded at 5 frames per second in low speed experiments and at 100 frames per second in high speed experiments.

ZMW experiments were essentially performed under the at same conditions as TIRF microscopy, except for the following details. 30S PICs immobilization concentration was increased to 200 nM. Upon washing off unbound complexes with the reaction buffer, the chip was overlaid with 20 μl of fresh polymix buffer supplemented with 1 μM IF2, 2.5 mM TSQ, 2.5 mM protocatechuic acid, and 1x protocatechuate-3,4-dioxygenase. Translation was started by delivering 20 μl mixture of 50S ribosomes, ternary complexes and EF-G with all reagents at 2x concentration. The buffer in the chip and delivery mix would mix upon solvent delivery. The resulting reagent concentration is indicated in text. Fluorescence was recorded at 10 frames per second. In ZMW experiments, illumination and delivery were delayed relative to the start of data acquisition. This delay was recorded and accounted for during data analysis. The 532 nm illumination was used at 0.35 μW/ μm^2^

### Simultaneous tracking of subunit movement and EF-G occupancy

EF-G mutant S73C was purified and labeled with Alexa-588 maleimide as previously described (Chen et al., 2013). Both Cy3 and Alexa-588 could be independently excited with the 532 nm laser but have distinct emission profiles. It makes possible simultaneous tracking of Cy3-Cy5 FRET and Alexa-588 labeled protein in the ZMW instrument. The experimental conditions were the same as above, except that the illumination power was increased to 0.7 μW/μm^2^. The 566/28, 610/60 and 658.5/37 nm spectral windows were used to acquire Cy3, Alexa588 and Cy5 fluorescence correspondingly (Chen et al., 2014).

## References

1. Kaledhonkar, S., Fu, Z., Caban, K., Li, W., Chen, B., Sun, M., Gonzalez, R.L., and Frank, J. (2019). Late steps in bacterial translation initiation visualized using time-resolved cryo-EM. Nature 570, 400–404.

2. Klimova, M., Senyushkina, T., Samatova, E., Peng, B.Z., Pearson, M., Peske, F., and Rodnina, M. V (2019). EF-G-induced ribosome sliding along the noncoding mRNA. Sci. Adv. 5, eaaw9049.

3. Choi, J., and Puglisi, J.D. (2017). Three tRNAs on the ribosome slow translation elongation. Proc. Natl. Acad. Sci. 114, 13691LP–13696.

4. Noller, H.F., Lancaster, L., Zhou, J., and Mohan, S. (2017). The ribosome moves: RNA mechanics and translocation. Nat. Struct. Mol. Biol. 24, 1021–1027.

5. Prabhakar, A., Capece, M.C., Petrov, A., Choi, J., and Puglisi, J.D. (2017). Posttermination Ribosome Intermediate Acts as the Gateway to Ribosome Recycling. Cell Rep. 20, 161–172.

6. Belardinelli, R., Sharma, H., Caliskan, N., Cunha, C.E., Peske, F., Wintermeyer, W., and Rodnina, M. V (2016). Choreography of molecular movements during ribosome progression along mRNA. Nat. Struct. Mol. Biol. 23, 342–348.

7. Navon, S.P., Kornberg, G., Chen, J., Schwartzman, T., Tsai, A., Puglisi, E.V., Puglisi, J.D., and Adir, N. (2016). Amino acid sequence repertoire of the bacterial proteome and the occurrence of untranslatable sequences. Proc. Natl. Acad. Sci. U. S. A. 113, 7166–7170.

8. Sharma, H., Adio, S., Senyushkina, T., Belardinelli, R., Peske, F., and Rodnina, M.V. (2016). Kinetics of Spontaneous and EF-G-Accelerated Rotation of Ribosomal Subunits. Cell Rep. 16, 2187–2196.

9. Sprink, T., Ramrath, D.J.F., Yamamoto, H., Yamamoto, K., Loerke, J., Ismer, J., Hildebrand, P.W., Scheerer, P., Bürger, J., Mielke, T., et al. (2016). Structures of ribosome-bound initiation factor 2 reveal the mechanism of subunit association. Sci. Adv. 2, e1501502.

10. Chen, J., Coakley, A., O’Connor, M., Petrov, A., O’Leary, S.E., Atkins, J.F., and Puglisi, J.D. (2015). Coupling of mRNA Structure Rearrangement to Ribosome Movement during Bypassing of Non-coding Regions. Cell 163, 1267–1280.

11. Ling, C., and Ermolenko, D.N. (2015). Initiation factor 2 stabilizes the ribosome in a semirotated conformation. Proc. Natl. Acad. Sci. 112, 15874LP–15879.

12. Chen, J., Dalal, R. V., Petrov, A.N., Tsai, A., O’Leary, S.E., Chapin, K., Cheng, J., Ewan, M., Hsiung, P.-L.P.-L., Lundquist, P., et al. (2014). High-throughput platform for realtime monitoring of biological processes by multicolor single-molecule fluorescence. Proc. Natl. Acad. Sci. 111, 664–669.

13. Johansson, M., Chen, J., Tsai, A., Kornberg, G., and Puglisi, J.D. (2014). Sequence-dependent elongation dynamics on macrolide-bound ribosomes. Cell Rep. 7, 1534–1546.

14. Qin, P., Yu, D., Zuo, X., and Cornish, P. V (2014). Structured mRNA induces the ribosome into a hyper-rotated state. EMBO Rep. 15, 185–190.

15. Tsai, A., Kornberg, G., Johansson, M., Chen, J., and Puglisi, J. (2014). The dynamics of SecM-induced translational stalling. Cell Rep. 7, 1521–1533.

16. Chen, J., Petrov, A., Tsai, A., O’Leary, S.E., and Puglisi, J.D. (2013). Coordinated conformational and compositional dynamics drive ribosome translocation. Nat Struct Mol Biol 20, 718–727.

17. Tsai, A., Uemura, S., Johansson, M., Puglisi, E.V., Marshall, R.A., Aitken, C.E., Korlach, J., Ehrenberg, M., and Puglisi, J.D. (2013). The impact of aminoglycosides on the dynamics of translation elongation. Cell Rep. 3, 497–508.

18. Chen, J., Tsai, A., O’Leary, S.E., Petrov, A., and Puglisi, J.D. (2012). Unraveling the dynamics of ribosome translocation. Curr. Opin. Struct. Biol. 22, 804–814.

19. Aitken, C.E., and Puglisi, J.D. (2010). Following the intersubunit conformation of the ribosome during translation in real time. Nat Struct Mol Biol 17, 793–800.

20. Fischer, N., Konevega, A.L., Wintermeyer, W., Rodnina, M. V., and Stark, H. (2010). Ribosome dynamics and tRNA movement by time-resolved electron cryomicroscopy. Nature 466, 329–333.

21. Marshall, R.A., Aitken, C.E., and Puglisi, J.D. (2009). GTP Hydrolysis by IF2 Guides Progression of the Ribosome into Elongation. Mol. Cell 35, 37–47.

22. Aitken, C.E., Marshall, R.A., and Puglisi, J.D. (2008). An Oxygen Scavenging System for Improvement of Dye Stability in Single-Molecule Fluorescence Experiments. Biophys. J. 94, 1826–1835.

23. Cornish, P. V, Ermolenko, D.N., Noller, H.F., and Ha, T. (2008). Spontaneous Intersubunit Rotation in Single Ribosomes. Mol. Cell 30, 578–588.

24. Johansson, M., Bouakaz, E., Lovmar, M., and Ehrenberg, M. (2008). The Kinetics of Ribosomal Peptidyl Transfer Revisited. Mol. Cell 30, 589–598.

25. Marshall, R.A., Dorywalska, M., and Puglisi, J.D. (2008). Irreversible chemical steps control intersubunit dynamics during translation. Proc. Natl. Acad. Sci. 105, 15364–15369.

26. Ermolenko, D.N., Majumdar, Z.K., Hickerson, R.P., Spiegel, P.C., Clegg, R.M., and Noller, H.F. (2007). Observation of Intersubunit Movement of the Ribosome in Solution Using FRET. J. Mol. Biol. 370, 530–540.

27. Horan, L.H., and Noller, H.F. (2007). Intersubunit movement is required for ribosomal translocation. Proc. Natl. Acad. Sci. U. S. A. 104, 4881–4885.

28. Dorywalska, M., Blanchard, S.C., Gonzalez Ruben L., J., Kim, H.D., Chu, S., and Puglisi, J.D. (2005). Site-specific labeling of the ribosome for single-molecule spectroscopy. Nucleic Acids Res. 33, 182–189.

29. Blanchard, S.C., Kim, H.D., Gonzalez, R.L., Puglisi, J.D., and Chu, S. (2004). tRNA dynamics on the ribosome during translation. Proc. Natl. Acad. Sci. U. S. A. 101, 12893–12898.

30. Katunin, V.I., Muth, G.W., Strobel, S.A., Wintermeyer, W., and Rodnina, M. V (2002). Important Contribution to Catalysis of Peptide Bond Formation by a Single Ionizing Group within the Ribosome. Mol. Cell 10, 339–346.

31. Frank, J., and Agrawal, R.K. (2000). A ratchet-like inter-subunit reorganization of the ribosome during translocation. Nature 406, 318–322.

32. Rodnina, M. V, Savelsbergh, A., Katunin, V.I., and Wintermeyer, W. (1997). Hydrolysis of GTP by elongation factor G drives tRNA movement on the ribosome. Nature 385, 37–41.

